# TF-MAPS: fast high-resolution functional and allosteric mapping of DNA-binding proteins

**DOI:** 10.1101/2025.10.20.683418

**Authors:** Xianghua Li, Ben Lehner

## Abstract

Transcription factors (TFs) bind specific DNA sequences to control gene expression. Modulating TF activity is of considerable therapeutic interest but very few TFs have been successfully drugged. TF DNA-binding interfaces are large and most TFs have no known allosteric ligands or allosteric sites. Here, we introduce TF-MAPs, a scalable and general platform to chart complete functional maps of sequence-specific DNA-binding proteins. We apply TF-MAPS to three human disease TFs: HNF1A, FOXG1 and FOXP1. Allostery in all three proteins is distance-dependent, anisotropic, and couples the DNA-binding interface to the protein surface. The allosteric surfaces of FOXG1 and FOXP1 are quite diverged, revealing a potential for protein-specific regulation in protein families. All three TFs have potentially druggable pockets where mutations have strong functional effects on binding, either within the binding interface or distant from it. Using TF-MAPS it should be possible to chart functional and allosteric maps for hundreds of human TFs and other DNA-binding proteins to understand DNA recognition and TF regulation, to mechanistically interpret clinical variants, and to guide the development of TF-modulating therapeutics.

## Introduction

The human genome encodes >1,500 transcription factors (TFs), sequence-specific DNA-binding proteins that regulate gene expression. The DNA-binding specificities of hundreds of TFs have been quantified alone ^1^ and in combination ^2^, as has their binding across the genome in multiple cell types ^3^. Many TFs are high-value therapeutic targets. Rare variants in 164 TFs cause 277 human genetic diseases (ClinVar 11 Nov 2023) and many TFs are also implicated in common diseases ^4–6^. In addition, at least 190 TFs are encoded by cancer driver genes ^4,7^. Of these, 87 are oncogenes and 60 are tumour suppressors ^8,9,10^.

Despite their causal role in hundreds of human diseases TFs are considered very challenging therapeutic targets. TFs are not enzymes and so have no catalytic sites to inhibit using small molecules. Moreover, they bind DNA through large and typically charged interfaces much larger than drug-sized small molecules ^11,12^. In addition, for many TFs such as tumour suppressor genes, the desired therapeutic outcome is activation, not inhibition ^13^. For destabilising missense variants, small molecule stabilisers (also known as correctors or pharmacochaperones) may present a viable therapeutic strategy, and correctors have been developed to bind and stabilise p53 ^14^. In addition, the serendipitous discovery of molecular glues that recruit ubiquitin ligases to degrade oncogenic TFs has led to extensive efforts to develop new TF degrader therapeutics ^15^. Overall, though, there are no licensed or experimental drugs against the vast majority of TFs mutated or implicated in human diseases ^11^.

One class of human TFs that has been very successfully drugged is nuclear receptors. Nuclear receptors are a family of 48 human TFs that are activated or inhibited by the binding of small molecules such as hormones. Examples include the estrogen, progesterone, androgen, and steroid hormone receptors. Ligands bind these TFs at a site distant to the DNA-binding interface and indirectly alter the binding of the receptors to other proteins, their nuclear localisation and DNA by causing changes in conformation and dynamics ^16^. This mechanism of regulation is an example of allostery – the long-range transfer of information between distant sites in proteins ^17^. Over a third of nuclear receptors have been successfully drugged, with nearly 200 licensed inhibitors and activators approved by regulatory agencies ^18^. Indeed, nuclear receptors are the third most successfully drugged protein family (after GPCRs and ion channels), demonstrating the huge potential for TF-targeting therapeutics^19^. A second family of human TFs with known allosteric ligands is the bHLH–PAS (basic helix–loop–helix–PER–ARNT–SIM) family and small molecule inhibitors have been successfully developed against Hypoxia-Inducible Factor 2α (HIF2α) ^20^, the aryl hydrocarbon receptor (AHR) ^21^, and the master circadian regulator brain and muscle Arnt-like 1 (BMAL1) ^22^.

Allostery is central to biological regulation and also to the efficacy of many licensed drugs ^23^. However, most proteins, including nearly all TFs, have no known allosteric sites. Indeed, the extent of allosteric communication in structurally diverse proteins is largely unknown ^24^. Recently, powerful approaches have been developed to systematically and comprehensively quantify allosteric communication in proteins using mutations, providing the first complete experimental allosteric maps of a small number of proteins and protein domains ^25–28^. The general approach involves quantifying the effects of mutations in all sites in a protein on a selectable activity—for example, its binding to another protein or its catalytic activity—and also quantifying the effects of the same mutations on the concentration of the folded protein. Combining the two selections—for example by fitting energy models to shallow double mutant data^25,29,30^—allows the effects of mutations on activity not caused by changes in the concentration of the folded protein to be quantified. By definition, if a mutation outside of an active site alters activity or binding, it must be via an indirect or allosteric mechanism ^25–27^.

These first complete experimental allosteric maps have revealed that allostery is surprisingly pervasive, with mutations in many sites distant from active sites altering binding or catalytic activity. Moreover, the experimental allosteric maps generated to date all share a characteristic distance-dependent decay of allosteric mutational effects away from the active site of the protein: mutations are, in general, more likely to have allosteric effects on activity if they are in residues close to the active site and the magnitude of allostery decays approximately exponentially with 3D distance in the protein structure ^25–28,31,32^. Allostery is, however, anisotropic, with mutations in particular sites having stronger allosteric effects than other mutations at the same distance from the active site. We refer to these sites as ‘allosteric hotspots’ ^28^. This spatial and functional mapping of allosteric effects provides a powerful framework for identifying therapeutically actionable sites.

For drug discovery, solvent accessible surface sites and pockets are of particular interest. Several of the published allosteric maps have been used to annotate potentially druggable surface pockets according to whether mutations in the pockets do or do not have strong allosteric effects. In both the Src kinase and KRAS, multiple known and novel pockets have been prioritised for therapeutic targeting because they are enriched for allosteric mutations ^26,27^.

Here, we present a general and fast experimental platform that extends allosteric mapping to DNA-binding proteins, including TFs. Experimentally, the strategy—that we term TF-MAPS (Transcription Factor-Mutational Allosteric Propagation Scan)—has two steps. First, we use a massively parallel selection experiment to quantify the effects of mutations on sequence-specific binding to DNA. Second, we use a pooled selection experiment to quantify the effects of the same mutations on the concentration of the folded protein in the same cell type. We then combine these two datasets to quantify changes in DNA binding that are not caused by changes in folded protein abundance.

We first use TF-MAPS to chart a comprehensive allosteric map of the POU-Homeodomain (POU_H_) DNA-binding domain (DBD) of Hepatocyte Nuclear Factor 1 homeobox Alpha (HNF1A). HNF1A is a master regulator of gene expression in the liver, pancreas, kidneys, and intestines, including regulation of glucose metabolism, lipid metabolism, and insulin secretion ^33^. Autosomal dominant variants in HNF1A cause Maturity-Onset Diabetes of the Young type 3 (MODY3), a condition characterised by reduced insulin secretion due to pancreatic β-cell dysfunction ^33^. In addition, common single-nucleotide polymorphisms in HNF1A are associated with increased risk of Type 2 Diabetes Mellitus (T2DM) ^34–39^. Beyond its role in metabolic disease, both somatic and germline variants in HNF1A cause hepatic adenoma and carcinoma ^40,41^, with ∼35% of hepatocellular adenoma cases attributed to inactivating mutations in HNF1A ^40,41^. There are, however, no licensed drugs targeting HNF1A. HNF1A binds DNA through two domains: POU_H_ and a POU-specific domain (POU_S_). Both domains have pathogenic variants in MODY3, but only the POU_H_ is frequently mutated in hepatocellular adenoma ^42^.

We next apply TF-MAPS to a distinct TF family and generate comparative functional maps for two forkhead box (FOX) TFS, FOXG1 and FOXP1. FOXG1 and FOXP1 have the largest number of known pathogenic variants among the 43 human forkhead family TFs ^43,44^. Both FOX TFs function during development, and mutations cause rare severe neurodevelopmental disorders called FOXG1 syndrome and FOXP1 syndrome, respectively ^45,46^. FOXP1 also plays an important role in immune regulation and has been characterised as a potential tumour suppressor in various malignancies ^47,48^.

The functional and allosteric maps of these three DBDs provide insights into DNA recognition, fold stability, and long-range energetic coupling. Allostery in all three TFs shows distance-dependent decay but also anisotropy, pinpointing allosteric hotspot residues and solvent-accessible sites and pockets that serve as potential targets for regulation and therapeutic intervention. All three proteins have potentially druggable pockets where mutations strongly perturb DNA-binding. For FOXG1 and FOXP1 our data prioritise a pocket in the binding interface, whereas for HNF1A the functional map prioritises a distal allosteric pocket. Comparison of the FOXG1 and FOXP1 maps suggests allosteric surfaces evolve quite rapidly in protein families. Using the TF-MAPS approach, it should be possible to produce functional and allosteric maps for hundreds of human TFs to understand DNA-recognition, mechanistically interpret clinical variants, and guide the development of allosteric therapeutics.

## Results

### TF-MAPS: rapid functional and allosteric mapping for DNA-binding proteins

We conceived a general approach called TF-MAPS (Transcription Factor-Mutational Allosteric Propagation Scan) to comprehensively quantify residue function and allostery in DNA-binding proteins. TF-MAPS aims to test if perturbations at every site in a protein directly or indirectly alter its binding to DNA. TF-MAPS uses mutations as perturbations, for example, introducing all 19 alternative amino acids (aa) at each residue. Extensions to insertion and deletion variants may enhance mapping in future implementations ^49^.

TF-MAPS has two experimental steps. First, the effects of all mutations in all sites on the protein’s ability to bind to a specific DNA sequence are quantified. Second, the effects of the same mutations on the protein’s folded cellular abundance are quantified. Integrating these two measurements then allows changes in binding beyond those caused by changes in concentration of the folded protein to be determined, providing a complete picture of how mutations throughout a protein quantitatively alter its ability to bind to DNA (Fig.1a).

One goal of TF-MAPS is to identify the residues contacting DNA that are most important for binding. In protein-protein interactions, such residues are referred to as ‘binding interface hotspots’ ^50,51^. Importantly, TF-MAPS also systematically evaluates whether mutations at all other sites alter DNA binding; by definition, any change in binding caused by a mutation in a residue that does not contact DNA must have an indirect or allosteric mechanism of action ^25–28,31,32,52^. The goal of TF-MAPS is not to identify or discriminate between alternative allosteric mechanisms (for example, conformational changes vs. changes in dynamics ^24^). Rather, the objective is to identify all sites where mutations have allosteric effects on DNA-binding and to quantify the strength of these effects.

To quantify DNA-binding, TF-MAPS employs a modified Omega bacterial one-hybrid (B1H) assay, which uses a fusion between the transcription factor of interest and a subunit of RNA polymerase ^53–55^, enabling the protein interaction with a target DNA sequence to regulate expression of a reporter gene (HIS3). The level of reporter gene expression directly influences the cellular growth rate, thereby providing a quantitative measure of DNA-binding activity (Fig.1b).

To quantify folded protein abundance in the same cellular context, TF-MAPS uses a spectinomycin tripartite sandwich (STS) assay ^56^. Insertion of the DBD into a split streptomycin-resistance protein links bacterial growth to TF folding, as only the folded protein confers antibiotic resistance ^52,56,57^.

The TF-MAPS approach is inspired by methods developed for producing allosteric maps of protein-protein interaction domains ^25,26,28^, kinases ^27^, and receptors ^32,52,58^. However, because TF-MAPS is conducted entirely in bacteria, it offers the advantages of speed, simplicity, and smaller volumes, making it more scalable.

### Functional map of HNF1A

We first applied TF-MAPS to an important human disease TF DBD, the POU-Homeodomain (POU_H_) from HNF1A (Fig.1c; Extended Data Fig.1a,b). Rare variants in the POU_H_ of HNF1A cause a form of monogenic diabetes, Maturity-Onset Diabetes of the Young type 3 (MODY3) ^33^. Somatic and germline variants in the POU_H_ domain also cause hepatocellular adenoma ^42^.

We used pooled DNA synthesis to construct a library containing all 19 aa substitutions as well as stop and synonymous variants in all sites in the DBD (residues 197 to 288), a total of 1748 missense variants, 350 synonymous variants, and 92 premature stop variants (Fig. 1d). We then quantified the ability of each of these HNF1A variants to bind to a consensus DNA motif using the B1H system, and quantified their effects on the concentration of folded HNF1A protein using STS (Fig. 1b,e). Replicate measurements for both assays were highly correlated (median Spearman’s ρ = 0.944 for B1H replicate transformations of the library and Spearman’s ρ = 0.938 for STS at three concentrations of spectinomycin 593, 445, and 334 μg/mL, p < 2.2e-16 for both) (Fig.1f; Extended Data Fig.1c-f). For both assays, enrichment scores were normalised using the mode of synonymous (normalised fitness = 0) and stop (normalised fitness = -1) variants (Fig.1g).

**Fig. 1:**
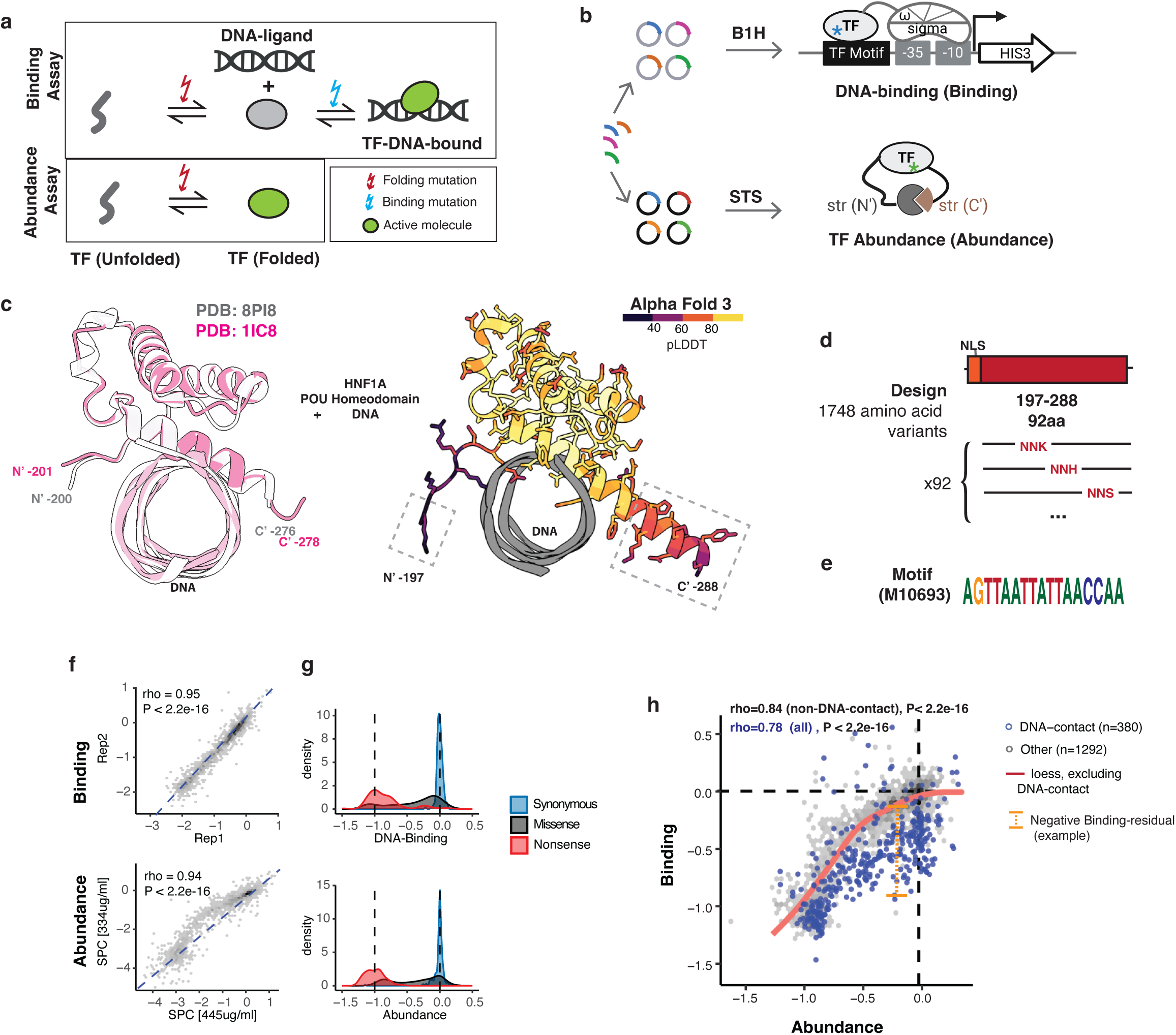
TF-MAPs. **a**, Schematic overview of mutational effects assessed via DNA-binding and protein abundance assays, illustrating their impact on transcription factor (TF) states. **b**, Reporter systems used in the experiments: bacterial one-hybrid (B1H) assay for DNA-binding activity, and a spectinomycin-based tripartite assay for protein abundance (SS). **c**, Structures of HNF1A POU_H_ bound to DNA: experimentally solved (left) and AlphaFold3-predicted (right). Dashed boxes highlight regions absent in the solved structure. **d**, Composition of the mutant library targeting HNF1A POU_H_ and flanking regions. NLS, nuclear localisation signal. Degenerate codons: N = A/T/G/C; K = G/T when WT is C or A; S = C/G when WT is A; H = A/C/T when WT is G. **e**, HNF1A binding motif used in the DNA-binding assay. **f**, Assay reproducibility and mutation distributions before scaling mutational effects between 0 and –1. DNA-binding assay shows correlation between biological replicates; abundance assay shows correlation across two conditions. **g**, Distribution of mutations across functional classes after normalisation (0 = wild-type; –1 = mode of nonsense mutations). **h**, Relationship between abundance and binding effects, with residuals computed from a loess fit on non-DNA-contacting residues.

Plotting the binding of each HNF1A variant against its abundance reveals a strong non-linear correlation (Spearman’s ρ = 0.78, p < 2.2e-16, Fig.1h), consistent with many changes in binding being caused by changes in abundance, as also observed for protein-protein interaction domains ^25,26,28^ and receptors ^32,52,59^. As expected, mutations in the hydrophobic core are very detrimental for abundance compared to mutations in other sites (median abundance = -0.82, Kruskal-Wallis chi-squared = 331, p < 2.2e-16) (Fig.2a, c, f, i, l; Extended Data Fig. 2a, b), with substitutions to proline most detrimental (median = -0.91), particularly in secondary structure elements (median = -0.95) (Fig. 2a, c, f, i, l; Extended Data Fig. 2c). Also as expected, mutations in the 20 residues that contact DNA (see Methods) are very detrimental for binding (median binding = -0.37 for all 20 residues and -0.45 for 12 residues in the structured helixes), often without causing large changes in abundance (Fig.2h-m).

**Fig. 2:**
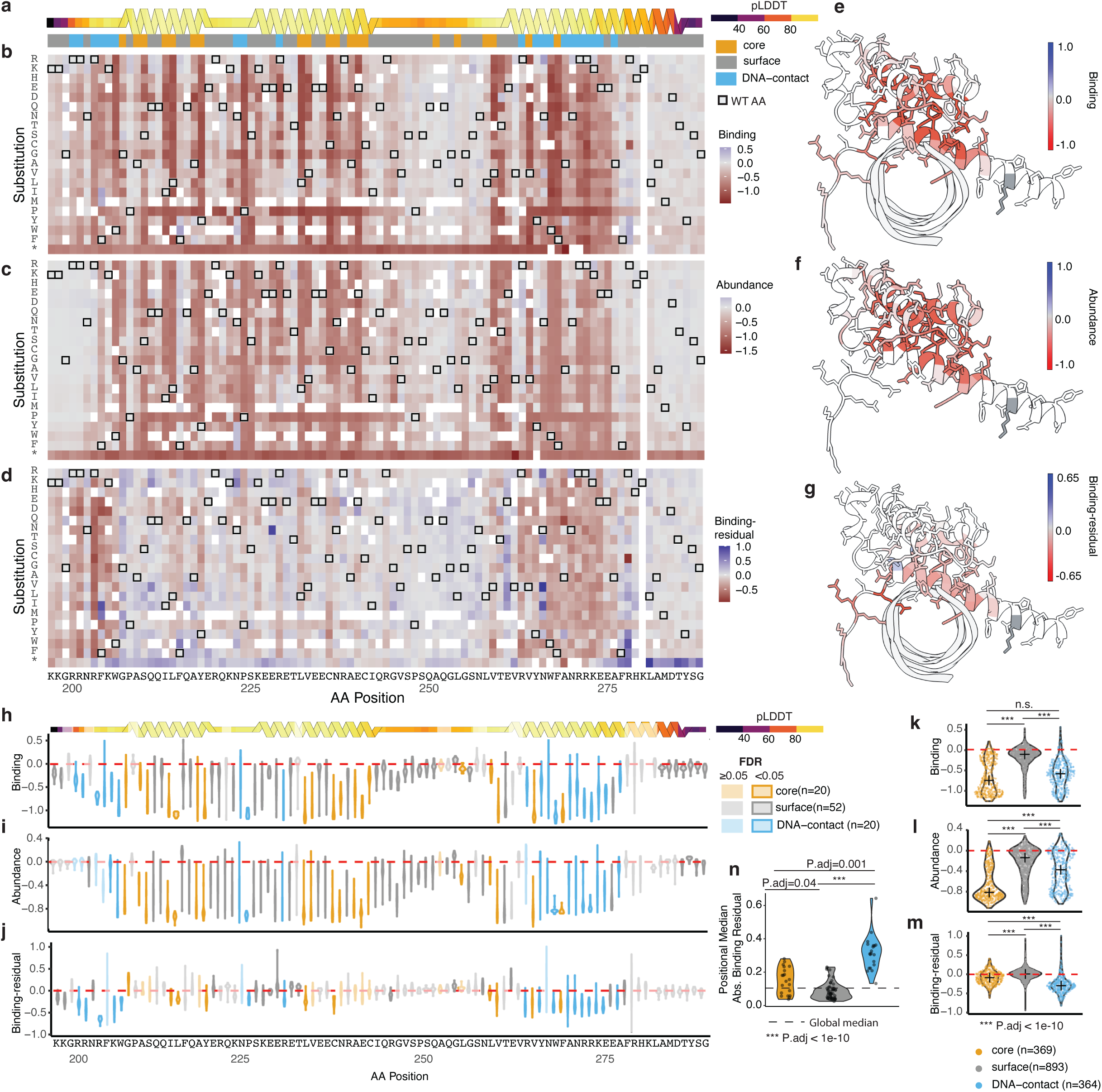
**a**, Structural annotation of residues based on the AlphaFold3-predicted DNA-bound structure. **b–d**, Heatmaps showing mutational effects from the DNA-binding assay (b), protein abundance assay (c), and binding-residuals (d), with missing values left blank. **e–g,** Median mutational effects per amino acid residue for DNA binding (e), abundance (f), and binding-residuals (g). The position without data is shown in dark grey. **h–j**, Violin plots of positional median effects for DNA binding (h), abundance (i), and binding-residuals (j). Statistical significance was assessed using one-way Wilcoxon tests followed by Benjamini–Hochberg false discovery rate correction. **k–m**, Comparison of mutational effects across three structural classes—core, surface, and DNA-contacting residues—for DNA binding (k), abundance (l), and binding-residuals (m). Statistical significance was assessed using Kruskal-Wallis tests followed by post-hoc Dunn’s tests with Bonferroni correction. **n**, Comparison of positional median absolute binding-residuals across the three structural classes. Kruskal–Wallis tests followed by post hoc Dunn’s tests were used to calculate P-values for panels k–n.

To quantify the impact of each mutation on binding beyond that caused by its measured abundance change, we used LOESS (Locally Estimated Scatterplot Smoothing) regression to infer the expected change in binding for each variant, given its folded abundance (see Methods) ^31,52^. For each mutation, the binding-residual between the measured and expected binding change quantifies its effect on binding beyond that caused by the measured change in folded abundance (Fig. 1h). These binding-residuals are well-distributed across the dynamic range of abundance measurements (Extended Data Fig. 2d).

### The HNF1A POU_H_ DNA interface

As expected, many mutations in the DNA-contacting residues have large binding-residuals (Fig. 2a, d, g, j, m; Extended Data Fig.3a). However, not all residues that contact DNA are equally important for binding (Fig. 2j, m). Overall, mutations in Loop 1 of the POU_H_ domain produce the strongest negative effects on DNA binding, followed by those in Helix 3 (Fig. 3g). In contrast, mutations in Loop 2 and Helix 2 have smaller effects, suggesting a hierarchy of functional importance among DNA-contacting regions (Fig. 3).

**Fig. 3:**
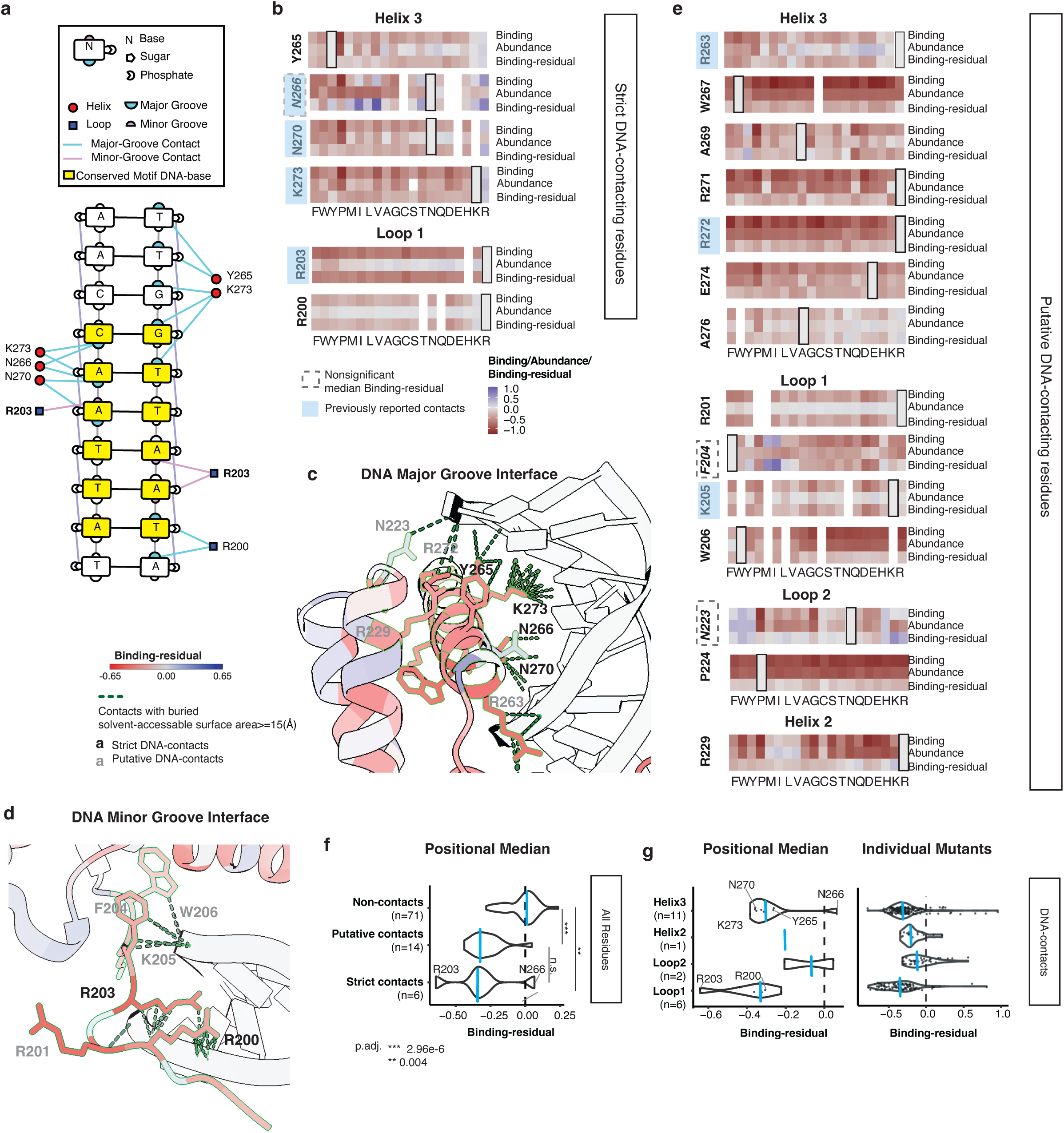
HNF1A binding interface. **a,** DNA-contact map generated with DNAproDB based on the AlphaFold3-predicted structure of the DNA-bound monomer. **b**, Mutational effects at residues identified as DNA contacts by DNAproDB (strict contact). **c, d**, DNA-bound structure of HNF1A POU_H_ highlighting broadly-defined DNA-contacting residues based on the ChimeraX “contacts” function at the major (c) and minor (d) grooves. Side chains of contacting residues are shown in atomic detail; the rest of the protein is depicted as a backbone cartoon. Dashed lines indicate protein–DNA contacts. **e**, Mutational effects at residues classified as putative DNA contacts (broader DNA-interface). **f, g**, Comparison of the mutational effects on binding-residuals: grouped by DNA-contact (f) and by DNA-contacting residues within different secondary structural elements (g).

DNAproDB ^60^ reports six residues (four in Helix 3: N266, N270, K273, Y264; and two in Loop1, R200 and R201) with base-specific contacts with DNA in the Alphafold3-predicted structure, which we refer to as strict contacts to differentiate them from putative contacts defined by changes in solvent-accessibility upon binding (see Methods) (Fig. 3a, Extended Data Fig. 3a). Several residues previously reported to contact DNA ^61,62^, but not identified by DNAproDB—including R263 and R272 within Helix 3, and R203 and K205 within Loop 1—also contribute significantly to DNA binding. Mutations in both direct and putative contacts similarly reduce binding-residuals (Fig. 3b-f).

Overall, eleven residues within Helix 3 contact DNA and mutations in ten of these impair DNA binding, whereas mutations in the remaining residue, N266, both enhance and impair binding (Fig. 3b). Helix 3 harbours the conserved WFXNXR motif (residues 267–272) that inserts into the major groove of DNA and constitutes the primary DNA-binding interface^63^ (Fig. 3c). Within the WFXNXR motif, all five DNA-contacting residues—W267, A269, N270, R271, and R272—exhibit strong negative binding-residuals upon mutation (Fig. 3b, c, e). Mutations in N266 increase (eight) and decrease (five) binding (absolute binding-residual ≥ 0.1), with substitutions to hydrophobic residues and arginine having particularly high binding-residuals (Fig. 3b). The most important positions for DNA binding beyond Helix 3 are R203 and K205 in the N-terminal linker region, which form extensive minor groove contacts^64,61^ and exhibit strong negative binding-residuals (Fig. 2j, 3b,d, e).

### Mutational evidence for an extended binding interface

Interestingly, Alphafold3^65^ predicts three additional DNA-contacting residues—R200, R201 and A276—not visible in the experimental HNF1A-DNA structures^61,64^. We used our data to evaluate these predictions (Extended Data Fig. 3). Residues 197–201 in the linker between the POU_S_ and POU_H_ domains (Loop 1) are unresolved in the experimental structures and form part of the nuclear localisation signal ^66^. Genetic variants in this region have been proposed to be pathogenic because they disrupt nuclear localisation ^66^. However, AlphaFold3 ^65^ modelling of the DNA-bound structure predicts that two residues (R200 and R201) directly contact DNA and our data show that mutations at these sites indeed strongly impair DNA-binding, suggesting direct interference with DNA binding as a pathogenic mechanism (Fig. 2a-j, Fig.3a, b, d, e; Extended Data 3f).

Residue A276 is resolved as part of Helix 3 in one DNA-bound structure (PDB: 8pi8) ^64^ but is located in a loop in another DNA-bound structure (PDB: 1ic8) ^61^ (Extended Data Fig. 3g). In neither structure does A276 make direct contact with DNA. However, in the AlphaFold3 model A276 is positioned within an extended Helix 3 and forms a backbone hydrophobic contact (Alanine’s Cβ with the methyl group of thymine’s C7 atom at a distance of 4.773Å) with the non-consensus region of the DNA ligand (Extended Data Fig. 3g). Consistent with this prediction, our data shows that mutations in A276 strongly reduce DNA-binding (Fig. 3e).

### Distance-dependent allosteric decay

We next considered residues outside of the DNA binding interface (Extended Data Fig. 4a-d). Plotting the absolute binding-residuals of mutations against the minimal 3D distance of each residue to the DNA reveals a striking relationship: the binding-residuals are strongest for mutations in the binding interface, but they are also stronger for mutations in residues closer to the interface, with the magnitude of the absolute binding-residuals decaying approximately exponentially with increasing distance from DNA (Fig. 4a).

**Fig. 4:**
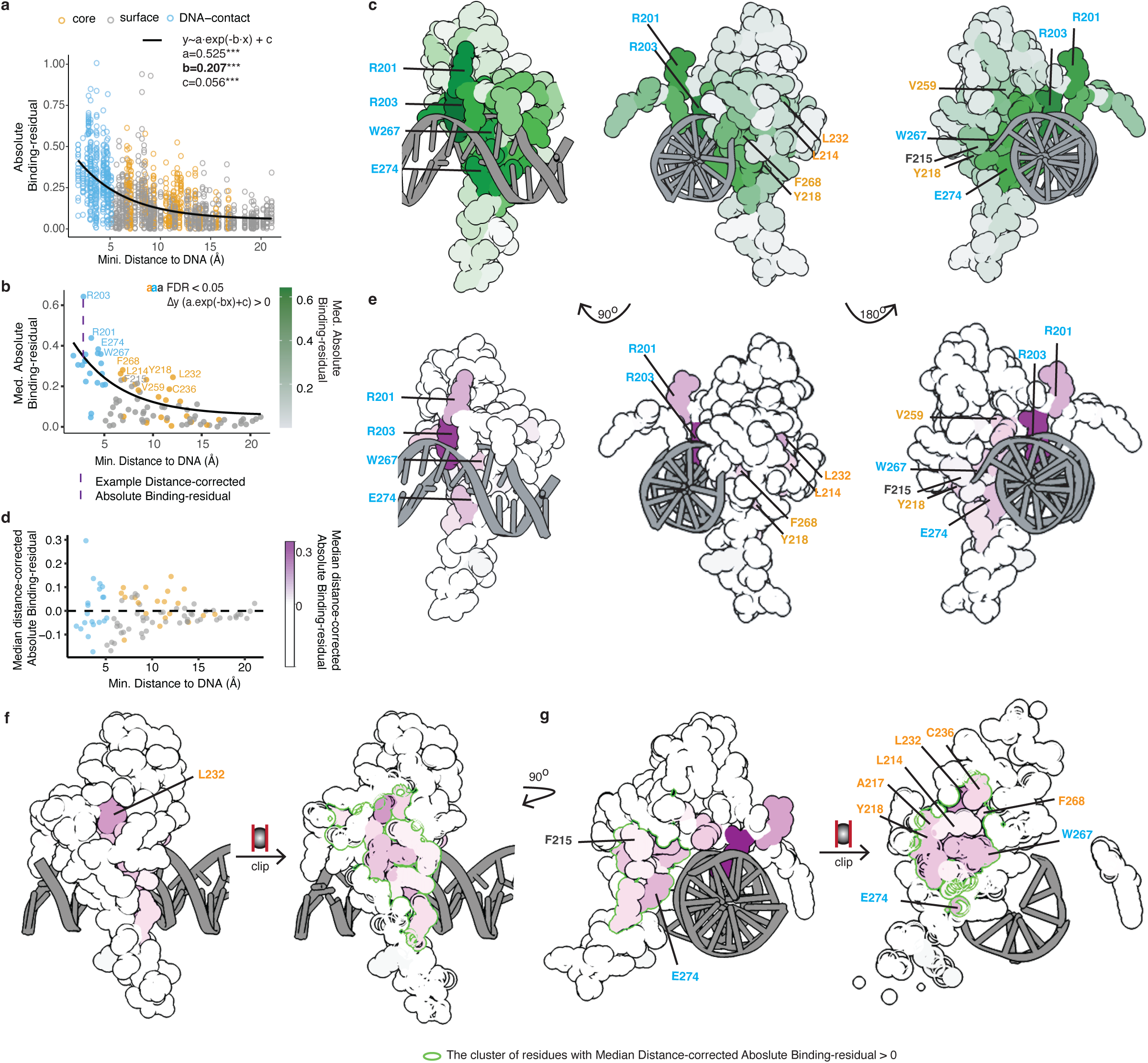
Allosteric regulation of DNA-binding in HNF1A. **a, b,** Positional decay plots of absolute binding-residuals for all mutations (a) and positional medians (b), coloured by structural annotation. **c,** Structural views of absolute binding-residuals from multiple perspectives, highlighting positions with significantly elevated residuals compared to decay-predicted values (FDR < 0.05, Wilcoxon test). **d,** Residuals of DNA-binding-residuals after fitting a decay curve based on distance to DNA. **e–g,** Structural views of allosteric anisotropy from different angles. Panels f and g highlight a cluster of residues with elevated distance-independent binding-residuals, shown from the perspective away from DNA (f) and from the side (g). The boundary of the cluster is outlined in green.

Plotting the median absolute binding-residual for mutations in each site further illustrates this (Fig. 4b), as does plotting the median binding-residuals on the structure of the DNA-bound protein (Fig. 4c; Extended Data Fig. 4e,f). This distance-dependent decay of allosteric effects is strikingly similar to that observed in protein-protein interaction domains ^25,26,28^, a hormone receptor ^32^, and an enzyme ^27^, suggesting it is likely to be a universal principle of protein biophysics.

### Allosteric anisotropy and hotspots

Although the allosteric effects of mutations on DNA-binding generally decay with distance, there is a wide dispersion of binding-residuals for different mutations (Fig. 4a) and residues (Fig. 4b) at each distance. To better characterise this variation in allostery across the protein, we first calculated the deviation of each site’s median absolute binding-residual from the value expected based on its distance to DNA (Fig. 4d). We then mapped these distance-corrected absolute binding-residuals onto the structure of the protein (Fig. 4e; Extended Data Fig. 4g).

This visualisation reveals that mutational effects on binding are particularly strong in a spatially clustered set of 16 residues located on the far side of helix 3 from the DNA-binding interface (Fig. 4f, g). These include partially buried residues A217, Y218 and L232 (rSASA 6-11%), and fully buried residues L214, V264 and F268 (rSASA=0). These connect via surface-exposed residues F215, E261 and R278 (rSASA>25%) to four DNA-contacting residues (W267, A269, R271, and E274) (Fig. 4e-g).

We define allosteric hotspots as positions outside the binding interface where mutations exhibit distance-corrected absolute binding-residuals significantly greater than zero (one-sample, one-sided Wilcoxon signed-rank test, BH-corrected FDR<0.05). In total, HNF1A has seven allosteric hotspots (Fig. 4b). When visualised on the HNF1A structure, all the identified hotspots localise within the interconnected cluster of 16 residues showing higher-than-expected binding-residuals (Fig. 4e-g). This cluster extends from the surface-exposed residue F215 (rSASA = 37%) through core residues L214 (rSASA = 0%), A217 (rSASA = 11%), Y218 (rSASA = 6%), L232 (rSASA = 11%), C236 (rSASA = 0%), and F268 (rSASA = 1%), and links to two DNA-contacting residues, W267 and E274, both of which also exhibit elevated distance-corrected absolute binding-residuals (Fig. 4f, g).

### HNF1A surface pockets

For regulation and drug discovery, solvent accessible surface sites and pockets are of particular interest. In total, 112 mutations in 11 surface sites of HNF1A have allosteric effects (FDR < 0.05 and absolute binding-residual ≥ 0.1), suggesting substantial potential for regulating HNF1A binding activity via interactions or perturbations of the surface. Structural pockets are of particular interest as potential small-molecule binding sites. Using the FPocket ^67^ algorithm, we identified six pockets in the HNF1A POU_H_ domain, including three distal to the DNA-binding interface (Pockets 1, 2, and 5) and three overlapping the interface (Pockets 3, 4 and 6) (Fig. 5a, Supplementary Table 2).

**Fig. 5:**
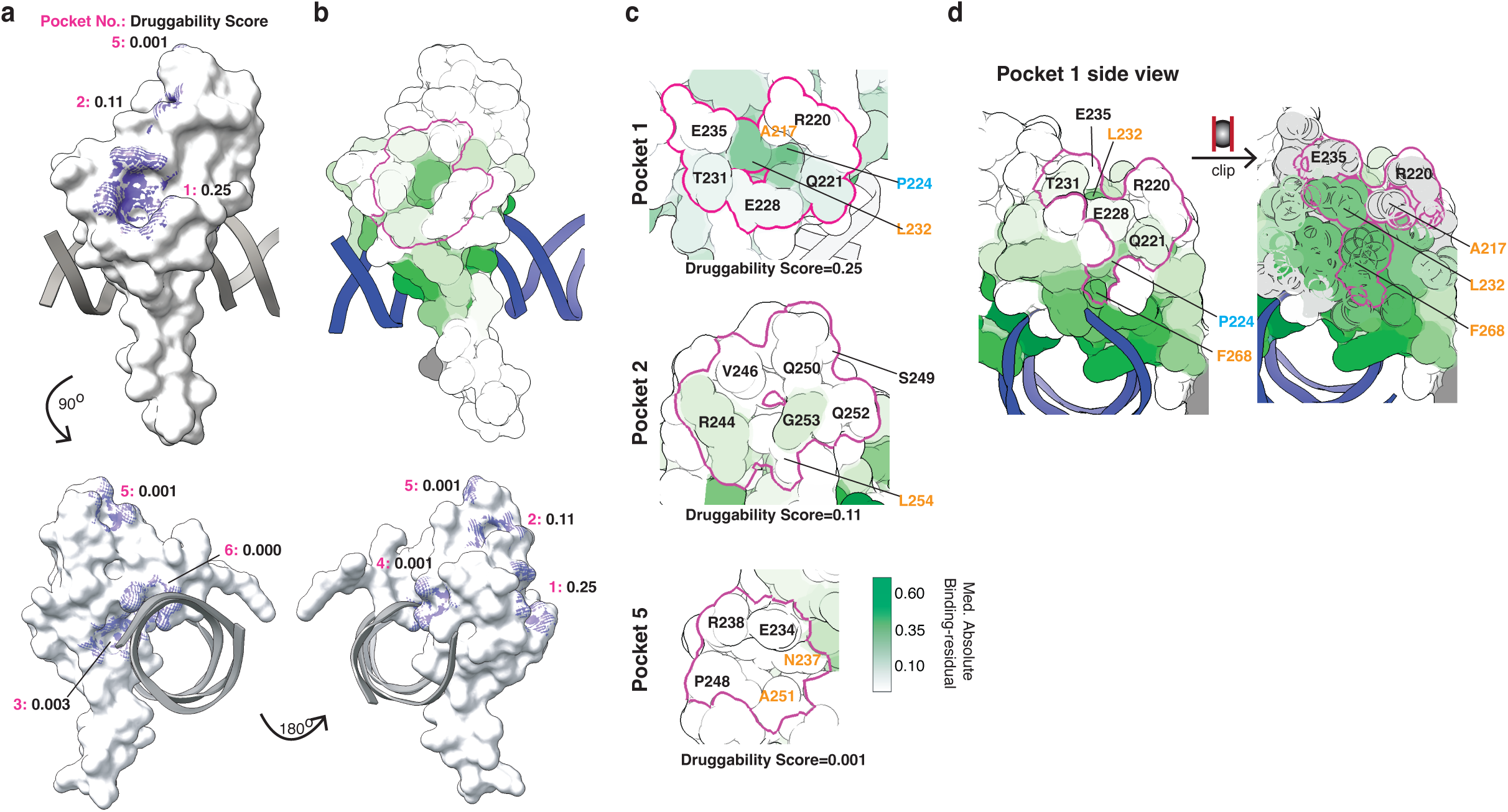
HNF1A pocket prioritisation. **a**, All pockets identified by the FPocket algorithm shown on the DNA-bound structure from multiple perspectives; pocket-forming residues are shaded. **b**, Pocket 1 residues viewed from the opposite side of DNA, encircled in magenta. Each residue is coloured by its positional median absolute binding-residual. **c**, Top views of three pockets located distal to DNA, with visible residues of the pockets labelled with the colour code corresponding ot the structural annotation of the residue. **d**, Side view of Pocket 1, showing all its constituent residues.

Our data show that mutations in all three pockets in the binding interface impair binding, but all three are also very shallow pockets predicted to have low druggability (Fig. 5a; Extended Data Fig. 5a-d, Supplementary Table 2). Of the three more distal pockets, mutations in Pocket 5 have little effect on binding. At least one substitution in each of the seven residues in Pocket 2 has a significant binding-residual, with mutations at two surface residues having a directional trend: most substitutions at R244 reduce binding, while most at G253 increase binding (Fig. 5c,

Extended Data Fig. 5f). However, Pocket 2 is quite shallow and has low predicted druggability. In contrast, Pocket 1 is predicted to have higher druggability and mutations in all five residues in the pocket affect binding. Indeed the pocket contains two allosteric hotspot residues, L232 and F268 (Fig. 4b, f, g, Fig. 5c-d, Extended Data Fig. 5e). Mutations in Pocket 1 residues both enhance and impair DNA-binding activity (Extended Data Fig. 5e), indicating potential for bidirectional modulation of HNF1A function.

The allosteric map therefore prioritises Pocket 1 as an allosterically active and drug-sized pocket on the surface of HNF1A POU_H_ (Supplementary Table 2).

### Mechanistic classification of pathogenic variants

Finally, we used the HNF1A TF-MAP to understand the mechanistic effects of pathogenic diabetes-causing variants (Fig. 6a). Among eighteen HNF1A variants annotated as pathogenic in ClinVar, sixteen reduce DNA binding in our assay (FDR < 0.05, Binding < -0.1) (Fig. 6a, b; Extended Data Fig. 6a, b). Fourteen localise to the DNA-binding interface, while one (G207D) reduces binding by destabilizing the protein, and another (E275V) acts via an allosteric mechanism (see Methods) (Fig. 6c; Extended Data Fig. 6). Two variants (R263H and V246L) show no detectable effects, suggesting a different mechanism-of-action (Fig. 6c; Extended Data Fig. 6a). Indeed, one of these variants, R263H, has previously been shown to disrupt nuclear localisation and reduces transactivation activity in a reporter assay ^68^. No functional studies have been reported for V246L, although V246 is solvent-accessible and part of allosteric Pocket 2 (Fig. 5b).

**Fig. 6:**
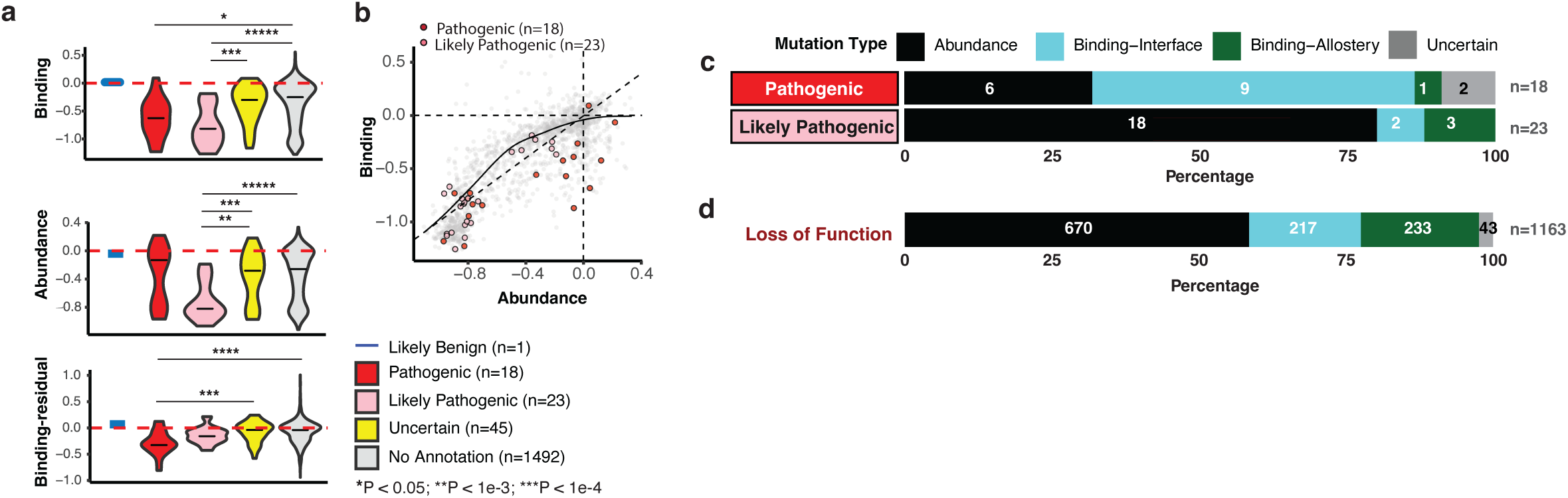
Mechanistic classification of HNF1A pathogenic variants. **a,** Functional scores compared across clinical groups. Statistical significance was assessed using Kruskal–Wallis tests followed by post hoc Dunn’s tests (*P* < 0.05; \*\**P* < 10⁻³; \*\*\**P* < 10⁻⁴). **b**, Two-dimensional binding–abundance landscape highlighting clinically annotated pathogenic and likely pathogenic variants. **c, d**, Classification of variants by biophysical category within pathogenic and likely pathogenic groups (c) and within loss-of-function mutations (d).

All 23 ClinVar variants annotated as ‘likely pathogenic’ reduce DNA binding in our assay (Fig. 6a, b). Six of these are in the DNA-binding interface (Extended Data Fig. 6a), while the remaining 17 act indirectly—14 by reducing protein abundance and three via allosteric mechanisms (Fig. 6c; Extended Data Fig. 6a).

In total our data identify a total of 1,163 missense variants that significantly reduce DNA binding (Extended Data Fig. 6b). Of these loss-of-function (LOF) variants, 57.6% act by destabilising the protein, while the remainder are almost equally divided between variants in the DNA-binding interface (18.7%) and variants that have allosteric mechanisms (20.0%) (Fig. 6d; Extended Data Fig. 6c).

This functional classification of known and predicted pathogenic variants is important for understanding disease mechanisms, for guiding therapeutic development, and for stratifying patients in clinical trials. Notably, since only a minority of the known pathogenic variants in HNF1A reduce binding via destabilisation, small molecule stabilisers ^69^ may have limited therapeutic benefit for most patients.

### Functional and allosteric maps of FOXG1 and FOXP1

To test TF-MAPs on a different DBD structural family and to compare the functional and allosteric maps of two homologous DBDs, we next applied TF-MAPS to two forkhead transcription factors, FOXG1 and FOXP1. Both proteins bind the canonical forkhead motif RYAAAYA ^47,70–73^ (R=G/A, Y=C/T) via Helix3 of the winged helix-turn-helix domain. Structural alignment of DNA-bound FOXG1 and FOXP1 DBD monomers shows high structural conservation (RMSD = 0.61Å; Fig. 7a). The sequences share 50% identity and 69% similarity with minimal gaps (6%) (Fig. 7a).

**Fig. 7:**
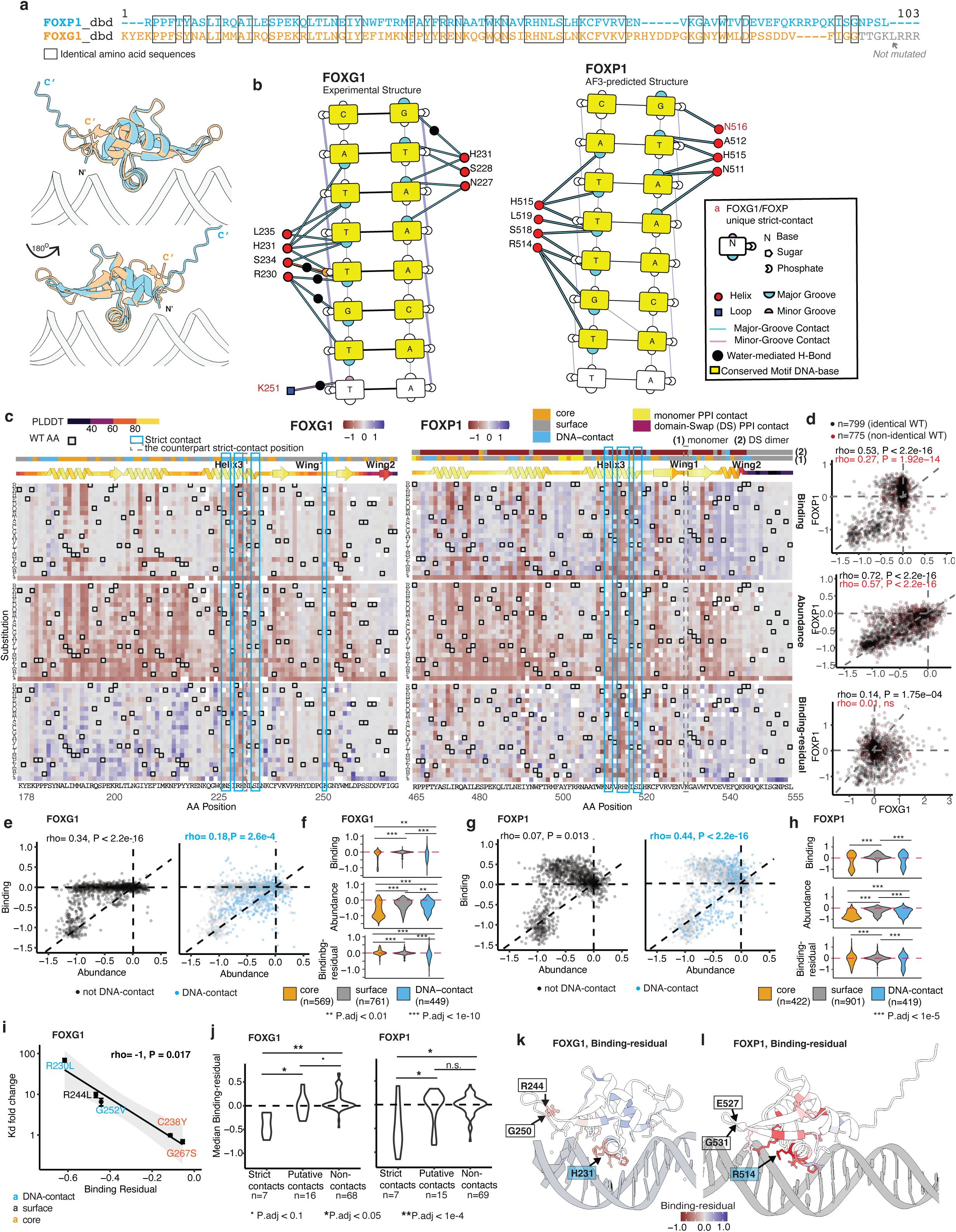
TF-MAPS for FOXG1 and FOXP1. **a,** Sequence and structural alignment of FOXG1 and FOXP1 forkhead DBDs. Grey residues in FOXG1 were not mutated in the study. **b**, DNA-contact map generated with DNAproDB based on the DNA-bound structure of FOXG1 (PDB:7CBY) and AlphaFold3-predicted structure of the DNA-bound FOXP1 monomer. **c**, Heatmaps showing mutational effects with missing values left blank. **d**, Correlations between matching variants’ binding, abundance and binding-residuals (from top to bottom) between FOXG1 and FOXP1. **e**, **g,** Relationship between abundance and binding effects for FOXG1 (e) and FOXP1 (g). **f, h,** Comparison of mutational effects across all structural classes for FOXG1 (f) and FOXP1(h). Statistical significance was assessed using Kruskal-Wallis tests followed by post-hoc Dunn’s tests with Bonferroni correction. P-values are shown only for statistically significant comparisons. **i**, Correlation between the published Kd (dissociation constant) values and the binding-residual of FOXG1. **j,** Comparison of mutational effects across all structural classes for FOXG1 and FOXP1. Statistical significance was assessed using Kruskal-Wallis tests followed by post-hoc Dunn’s tests with Bonferroni correction. P-values are shown only for statistically significant comparisons. **k, l,** FOXG1 (k) and FOXP1 binding-residuals (l). White-boxed positions indicate residues within Wing 1 that show large absolute binding residuals but do not contact DNA. For comparison, the DNA-contacting residue with the largest binding residual is highlighted with a coloured box.

We constructed variant libraries encoding all 19 amino acid substitutions, stop codons, and synonymous variants at each residue within the forkhead DNA-binding domains of FOXG1 (91 amino acids; residues 178–268 mutated out of 178-276, UniProt ID: P55316) and FOXP1 (91 amino acids; residues 465–555, UniProt ID: Q9H334) (Supplementary Table 1) and quantified sequence-specific binding to DNA using the B1H assay and folded protein abundance using the STS assay (See Methods). Mutational effects were highly reproducible across biological replicates in the B1H assay (median Spearman’s ρ = 0.966 for FOXG1, 0.979 for FOXP1, p < 2.2e-16) and across three consecutive drug concentrations in the STS assay (median Spearman’s ρ = 0.859 for FOXG1, 0.898 for FOXP1, p < 2.2e-16) (Extended Data Fig. 7a-i).

Mutational effects on abundance are well correlated between the two proteins (Spearman’s ρ = 0.72 for aligned identical wild-type residues and 0.57 for different wild-type residues, p < 2.2e-16) but less so than replicate measurements for each protein alone (Fig. 7c, d; Extended Data Fig. 7f, g; Extended Data Fig. 8a, b). Mutational effects on binding are less well correlated (Spearman’s ρ = 0.53 for aligned identical wild-type residues and 0.27 for different wild-type residues, p < 2.2e-13) (Fig. 7d; Extended Data Fig. 7b, 8; Extended Data Fig. 8).

Plotting DNA-binding against abundance for all mutations shows that, as for HNF1A, many changes in binding can be accounted for by changes in abundance (Fig. 7e-hi). Fitting a LOESS curve to the binding-abundance measurements allows a binding-residual to be calculated for each mutation (last rows in Fig. 7c). These binding residuals correlate very well with independently measured *in vitro* dissociation constants (K_d_) for FOXG1 ^73^ (Spearman’s ρ = –1, p = 0.017, n=5) (Fig. 7i; Extended Data Fig. 7j). For both proteins, the binding-residuals are large for mutations in the binding interface and in particular for mutations in Helix 3, which inserts into the DNA major groove (Fig. 7b, c, k, l). Across all identical residues, there is a small but significant correlation in binding-residuals between the two proteins (Spearman’s ρ = 0.14, p = 1.75e-4), but no correlation for mutations in non-identical wild-type residues (Spearman’s ρ = 0.01, n.s.) (Fig. 7d; Extended Data Fig.8b).

Mutations in residues identified by DNAproDB^60^ as direct DNA contacts have strong effects on binding, whereas most mutations in other DNA-interface residues—classified as putative contacts based on changes in solvent accessibility upon DNA-binding—do not (Fig. 7j). This contrasts with HNF1A, where no difference is observed between strict and putative contacts in their effects on DNA binding (Fig. 3f), indicating that in FOXG1 and FOXP1 only key interacting residues play a major role in mediating DNA binding.

Interestingly we observe strong binding-residuals for several positions within the Wing 1 region of both FOXG1 and FOXP1, despite no evidence for direct DNA contact in either experimental or predicted structures (Fig. 7k, l; Extended Data Fig. 7a). Mutations with large binding-residuals include those at an aligned glycine (G250 in FOXG1 and G531 in FOXP1), with median binding-residuals of -0.31 and 0.22, respectively (Fig. 7k, l). Additionally, mutations in nearby charged residues (FOXG1 R244 and an oppositely charged residue in FOXP1, E527) also have large residuals (-0.46 and -0.26, respectively) (Fig. 7a, k, l). Although these charged side chains do not directly contact DNA in the experimental structures of FOXG1 and FOXP1 ^73–75^, the Wing 1 region of another forkhead DBD–FOXD3–has been suggested to be flexible and capable of forming dynamic interactions with DNA ^76^. The mutational effects suggest this may also be the case in both FOXG1 and FOXP1. Alternatively, they may have an allosteric mechanism of action.

### Conserved distance-dependent allosteric decay and anisotropy

Mapping the absolute binding-residuals of all FOXG1 and FOXP1 missense mutations against their spatial distance to the DNA reveals the characteristic distance-dependent allosteric decay, consistent with the pattern observed in HNF1A (Fig. 8a and 4a, b). Besides this distance - dependent decay, both FOXG1 and FOXP1 exhibit notable dispersion of absolute binding-residuals at each distance (Fig. 8b, c, f), indicative of functional and allosteric anisotropy—where mutations in certain residues exert disproportionately strong effects, as observed in HFN1A.

**Fig. 8:**
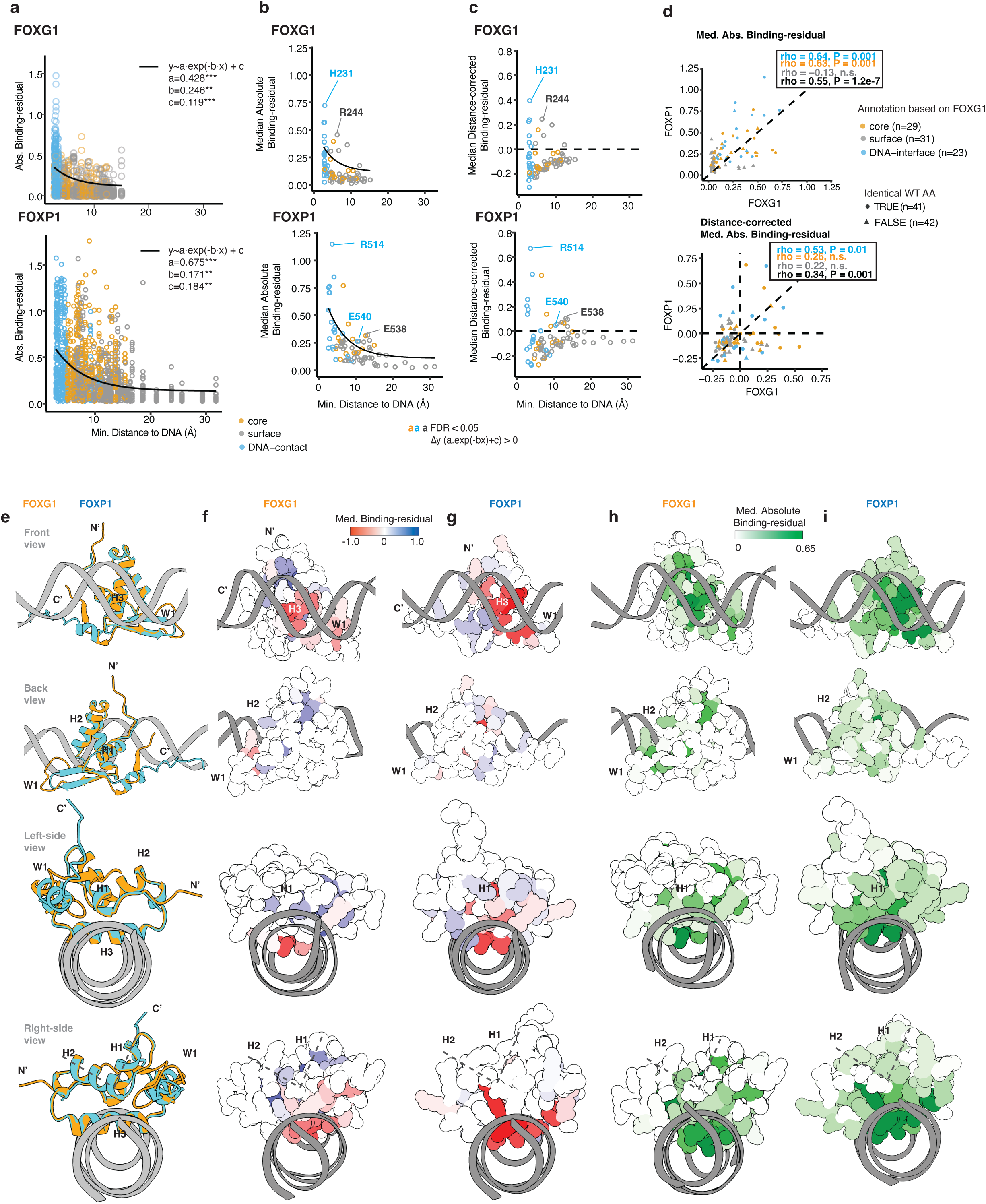
Binding interface and allosteric decay for FOXG1 and FOXP1. **a, b,** Positional decay plots of absolute binding-residuals for all mutations (a) and for position-wise median. **c**, Distance-corrected binding-residuals after fitting a decay curve based on distance to DNA, highlighting positions with significantly elevated residuals compared to decay-predicted values (FDR < 0.05, one-sample Wilcoxon signed-rank tests followed by BH-correction). **d,** Correlations between matching positions’ median absolute binding-residual and distance-corrected absolute binding-residual scores between FOXG1 and FOXP1. **e-i,** Structural views of aligned FOXG1 and FOXP1 (e); median binding-residuals of FOXG1(f) and FOXP1 (g); and median absolute binding-residuals of FOXG1 (h) and FOXP1 (i) from multiple perspectives.

The dispersion—quantified as distance-corrected absolute binding-residuals—is correlated between the two proteins (Spearman’s ρ = 0.34, p = 0.001) (Fig. 8d, Extended Data Fig. 8b). This correlation is observed for mutations in DNA-contacting residues (Spearman’s ρ = 0.53, p = 0.01) and for mutations at identical amino acids outside of the DNA-contacting residues (Spearman’s ρ = 0.45, p = 0.003) while no significant correlation is observed for sites with non-identical amino acids (Spearman’s ρ ≥ 0.22, n.s.; Extended Data Fig. 8b-d).

The allosteric anisotropy between the two domains is thus partially conserved, but notable differences in allosteric behaviour are observed, particularly at sites that are not conserved at the sequence level.

### FOXG1 and FOXP1 have diverged allosteric surfaces

We next compared the effects of surface mutations in the two homologous DBDs, visualised across four orientations relative to the DNA interface (Fig. 8e-i). In the front view, mutations in Helix 3 (H3), which contacts the major groove, have strong and conserved negative binding-residuals in both proteins (first row, Fig. 8e-g). Mutations in Wing 1 (W1) also have negative residuals in both proteins. In contrast, additional N- and C-terminal regions facing the DNA backbone have positive residuals for both DBDs (Fig. 8f, g).

From the back view, mutational effects are more divergent. FOXG1 exhibits stronger and more concentrated allosteric mutational effects, particularly along the left side of the protein’s back surface. FOXP1 shows milder and more diffusely distributed effects. (second row, Fig. 8e-g).

From the left-side view, where both the N- and C-termini face forward, the C-terminal end of H3 shows much stronger binding-residuals in FOXP1 compared to the corresponding positions in FOXG1 (third row, Fig. 8e-g). On the opposite side (right-side view), the region encompassing H1 and H2 exhibits the greatest differences: several residues in FOXG1 display positive median binding-residuals, whereas those in FOXP1 are close to zero. However, the absolute binding-residuals in FOXP1 are markedly higher in this region, indicating that mutations produce both positive and negative effects, yielding median binding-residuals near zero but elevated median absolute values (last row, Fig. 8e-i).

Overall, FOXP1 harbours a greater number of mutations with pronounced allosteric effects, as reflected by the absolute binding-residuals and the slower decay of binding-residuals with distance from the DNA (decay rate: 0.246 for FOXG1 and 0.171 for FOXP1) (Fig. 8a, h, i).

The effects of mutations in surface residues are not correlated in the two proteins (Spearman’s ρ = –0.02 and –0.13, both n.s. for median binding-residuals and median absolute binding-residuals; Fig. 8d; Extended Data Fig. 8c) compared to mutations in the core (Spearman’s ρ = -0.34, p = 0.046; absolute residuals Spearman’s ρ = 0.63, p = 0.001) and in DNA-contacting residues (Spearman’s ρ = 0.45, p = 0.03; absolute residuals Spearman’s ρ = 0.64, p = 0.001) (Fig. 8d; Extended Data Fig. 8c). This divergence in allostery in the surface sites in structurally homologous proteins has also been observed in PDZ domains ^28^.

### FOXG1 and FOXP1 pockets

Using FPocket ^67^ we identified three structural pockets in FOXG1 and eight in FOXP1 (see Methods) (Fig.9a, b; Supplementary Table 3 and Supplementary Table 4). All three FOXG1 pockets are located near the DNA and are structurally conserved in FOXP1. FOXP1 also has an additional five distal pockets not detected in FOXG1 (Fig. 9a, b; Extended Data Fig. 9c). Most of these pockets are shallow and have low predicted druggability (Fig.9a, b; Supplementary Table 3 and Supplementary Table 4), and in both proteins the pocket with the highest predicted druggability is Pocket 1. Pocket 1 is located in the binding interface and many mutations in the pocket strongly impair DNA binding, identifying it as a potentially druggable hotspot required for DNA-binding (Fig. 9a-e; Extended Data Fig. 9a-c).

**Fig. 9:**
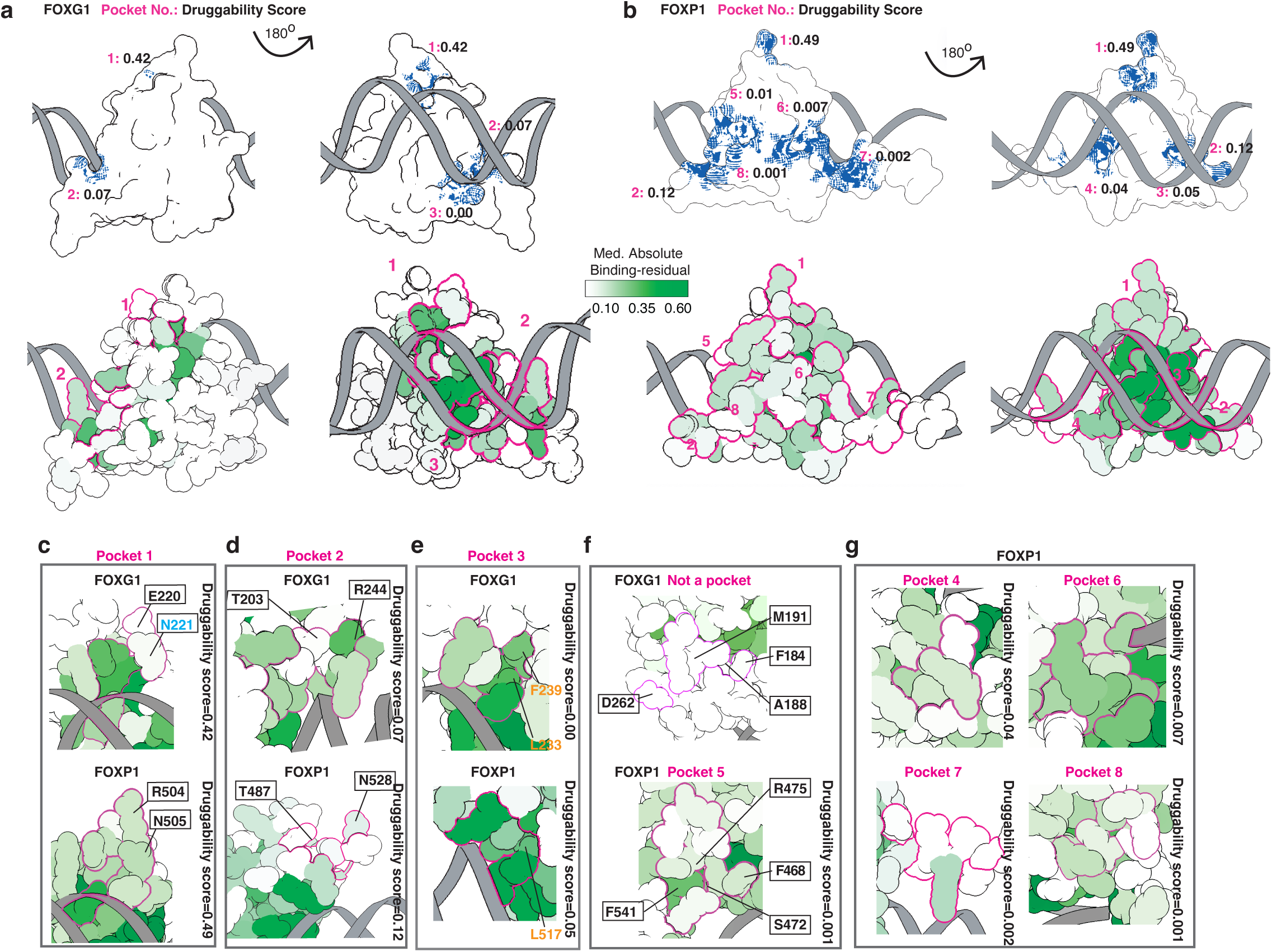
Allosteric surfaces and pockets of FOXG1 and FOXP1. **a, b,** All pockets identified by the FPocket algorithm shown on the DNA-bound structure for FOXG1(a) and FOXP1 (b); pocket-forming residues are shaded. **c-g**, Pocket-centric view, with each structurally aligned pocket pairs FOXG1 and FOXP1 Pocket 1 (c), FOXG1 and FOXP1 Pocket 2 (d), FOXG1 and FOXP1 Pocket 3, (e) and FOXP1 Pocket 5 with the corresponding positions in FOXG1 (f). The unmatched pockets of FOXP1 are shown together (g). Pockets are marked in magenta and individual residues coloured by their positional median absolute DNA-binding-residuals.

Mutations in the distal pockets of FOXP1 strongly affect DNA binding, suggesting the pockets are allosterically active (Fig. 9f, g; Extended Data Fig. 9a, d, e), with mutations in Pocket 6 having particularly strong allosteric effects (Fig. 9g; Extended Data Fig. 9e). However, targeting these shallow pockets is likely to be challenging because they are shallow with low predicted druggability (Fig. 9b; Supplementary Table 4).

In contrast to HNF1A, the functional maps therefore identify a binding interface hotspot pocket in FOXG1 and FOXP1 and prioritise it as the top pocket to target therapeutically with small molecules.

## Discussion

We have presented here a general and fast method—TF-MAPS—that uses mutations to produce complete functional and allosteric maps for DNA-binding proteins. TF-MAPS allows annotation of binding interfaces, identifies variant effect mechanisms, and produces global maps of allosteric regulation in DNA-binding proteins.

The functional maps of HNF1A, FOXG1 and FOXP1 provide important insights. First, all three proteins are allosterically active and show distance-dependent allosteric decay. Such decay is also observed in protein interaction domains ^25,26,28^, and an enzyme ^27^, suggesting it is a general principle of protein biophysics. Second, allostery in all three proteins is anisotropic, with a set of allosteric hotspot residues with unusually strong effects on binding when mutated. Third, the surface allosteric surfaces of FOXG1 and FOXP1 are divergent, suggesting each member of a TF family may have unique accessible allosteric sites to target for specific regulation ^28^. Fourth, the maps identify the DNA-contacting residues most important for DNA binding and provide experimental support for an extended DNA-binding interface for HNF1A involving a dynamic loop predicted by Alphafold3 and not observed in X-ray structures. Fifth, the maps provide reference atlases for clinical genetics, classifying the mechanisms-of-action for pathogenic variants. For example, considering loss-of-function variants in HNF1A, approximately half destabilize the protein, one quarter are in the binding interface, and one quarter have an allosteric mechanism-of-action. Sixth, the maps identify functionally important surface pockets to potentially therapeutically target. For FOXG1 and FOXP1 we suggest that a potentially druggable pocket in the DNA binding interface is the most promising pocket to target with small molecules, whereas in HNF1A a potentially druggable distal allosteric pocket exists where mutations have large effects on binding.

The current TF-MAPS approach has several important limitations, primarily that measurements are performed in bacteria on DNA that lacks the human chromatin context. Future work will be needed to assess how chromatinization influences mutational effects and allosteric behaviour ^77,78^. In addition, allosteric maps of KRAS binding to six different partners via overlapping interfaces are strongly correlated but not identical, suggesting target-specific allostery ^26^. Similarly, mutations can allosterically alter the peptide-binding specificity of a PDZ domain ^79,80^, and it will be interesting to quantify the extent to which allosteric effects are DNA sequence - specific for TFs. In addition, our allosteric map only covers the DBD and DBD-proximal regions. In future work, it will be important to extend functional and allosteric mapping to non-DBDs and the often large intrinsically disordered regions of human TFs ^81,82^.

A key strength of the TF-MAPS approach is its generalisability and scalability. With binding preferences available for hundreds of human TFs ^4^, we believe functional and allosteric mapping can be extended to a very large number of TFs relevant to both medicine and biotechnology.

Structure-based approaches can identify pockets in proteins, but they do not determine whether perturbation of these pockets has functional consequences. Mutational approaches such as TF-MAPS therefore build on structural biology to functionally annotate proteins, including building functional maps of protein pockets and surfaces. Challenging the view that TFs are undruggable proteins, all three TFs studied here have drug-sized pockets where mutations have strong effects on DNA-binding. The TF-MAPS approach is general and requires no specialist equipment. It can therefore be applied by many groups to the hundreds of different TFs implicated in human disease and important for biotechnology. Functional and allosteric maps of hundreds of human TFs and other DNA-binding proteins will provide a rich resource to understand, predict and engineer DNA recognition and TF regulation. The maps will also provide mechanistic atlases of variant effects to interpret germline and somatic clinical variants. Most importantly, however, we believe the approach will allow the identification of pockets and surface sites where perturbations have large effects on DNA binding. If these hotspot pockets are more widespread on TFs than is currently appreciated—as our data suggest they will be—it may be possible to transform TFs into successfully drugged targets, as has been achieved for other ‘undruggable’ protein families ^83^.

## Methods

### Variant Library Design and Preparation

All codon-randomised oligo pools (NNK, NNS, or NNH) were ordered from IDT to substitute each codon with variants encoding 19 alternative amino acids and stop codons. Here, “N” represents all four possible nucleotide bases. Depending on the wild-type nucleotide at the wobble position, we used different degenerate bases to reduce the occurrence of wild-type sequences and minimise the number of stop codons at each codon position. Specifically, we generally used the degenerate base K (G/T) to reduce stop codon frequency. When the wobble base is T, we used S (C/G) for substitution, and when the wild-type wobble base is G, we used H (A/C/T) to diversify the codon pool to minimise the occurrences of the wild-type nucleotide sequences.

For the 3′ primer oligo pools, the degenerate codon mixes were embedded in the reverse complement of the desired codon. For example, the reverse complement of NNK is MNN where M=C/A.

For HNF1A, FOXG1, and FOXP1, which have DNA-binding-domain (DBD) sizes <300 bp (100 amino acids), mutants were incorporated via PCR using oligo pools either as templates, 5′ primers (20–60 bp), or 3′ primers (20–60 bp). All 5′ ends of the forward and reverse primers, as well as the corresponding oligo pools for these three genes, shared constant regions to facilitate restriction enzyme digestion for downstream cloning (forward primer: cacccccccatggtacc; reverse primer: gcctttttctagtctctaga). Each PCR reaction was performed using NEB Q5 Hot-start polymerase for 10 cycles, followed by ExoSAP treatment to remove single-stranded oligos and purification using the MinElute kit.

All four TF DBD wild-type sequences were PCR-amplified from the MORF library (Addgene cat. no: 192821). Motif sequences were ordered as oligos from Sigma Aldrich, flanked by constant regions containing EcoRI and NotI sites for cloning into the pH3U3-zif268 (omega) backbone (Addgene cat. no: 18046).

The mutated region of HNF1A corresponds to amino acid positions 197–288 (UniProt ID: P20823) and PCR-amplified from the MORF library. The motif for HNF1A was obtained from CIS-BP (Motif ID: M10693; sequence: “AGTTAATTATTAACCAA”). Details of all the transcription factor DBDs in the study and their corresponding motifs are provided in Supplementary Table 1.

### Plasmid Library Preparation

To clone TF DBDs into B1H assay vector, we used the vector backbone of pB1H2w2-zif268 (Addgene cat.no: 18045), between the two restriction enzyme sites KpnI and Xba1 via digestion & T4 ligation following the manufacturer’s protocol. The binding motif was cloned into pH3U3 - zif268 (omega) (Addgene cat.no: 18046), between the two restriction enzyme sites NotI and EcoR1, using HiFi ligation following the manufacturer’s protocol of single-strand-double-strand ligation. For abundance assay, TF DBDs were cloned into plasmid backbone PTB182, which was acquired from the Bardwell lab^56^ and modified for flexible cloning in the Lehner lab.

In detail, each separate PCR reaction was performed for 10 cycles, followed by purification and cloning into plasmid backbones using XbaI and KpnI restriction sites. The PCR products were inserted into the XbaI–KpnI linearised vector pB1H2w2-zif268 ^54^, which was digested with the same restriction enzymes. Additionally, the PTB182 vector was linearised via PCR by introducing XbaI and KpnI sites between the two GS linkers, followed by enzymatic digestion and purification to prepare it for ligation.

T4 ligation was followed by transformation into NEB5α competent cells according to the manufacturer’s instructions. After recovery in 1 mL SOC, a 1 µL aliquot was diluted into 50 µL LB and plated on LB + Ampicillin plates, while the remainder was inoculated into 10 mL LB + Ampicillin for overnight culture. Colony counts from the plates were used to estimate the number of independent transformants. Libraries with at least 20-fold higher transformant numbers than variant counts were considered to have sufficient coverage. Plasmids were then prepared from the liquid culture, and the three same-gene mutant plasmid libraries were combined in equimolar amounts.

### DNA binding selections

The Bacterial One-Hybrid (B1H) assay was adapted from Meng and Wolfe ^53^, with key modification of using liquid rather than plate-based selection. USO hisB– pyrF– rpoZ– cells (Addgene cat. no: 18044) were made electrocompetent in-house and first transformed with motif-carrying plasmids. A colony with a validated plasmid was expanded, made electrocompetent again, and transformed with 100 ng of the library plasmid. An aliquot of recovery media was plated on 2xYT + carbenicillin + kanamycin to estimate transformant numbers; the remainder was cultured in 25 ml 2xYT + carbenicillin + kanamycin for each biological replicate.

After overnight growth (achieving >100x transformant coverage), 5 ml of culture was washed five times in water, incubated for 1 h at 37°C in NM media (no 3-AT), and finally inoculated into NM media with carbenicillin, kanamycin, IPTG as in the original protocol, and 1 mM 3-AT. Cultures were shaken at 37°C for 24 h, with aliquots measured in 96-well plates (3 technical replicates) for real-time OD monitoring, including WT and negative controls. The remaining wash media was spun to collect input cell pellets for -20°C storage.

After 24 h, output cell pellets were collected and plasmid-prepped along with input samples.

### Abundance selections

For the Spectinomycin Tripartite Sandwich Assay^56^, plasmids were electroporated into NEB 5-alpha electrocompetent cells. As with B1H, aliquots were plated to estimate transformant numbers; the rest was cultured in 25 ml LB + ampicillin. After confirming >100x variant coverage, 50 μl of overnight culture was seeded into LB + ampicillin with serial spectinomycin concentrations (diluted from 1000 mg/μl stock at 0.75x intervals). Three biological replicates were assigned to three consecutive concentrations across 21 dilutions. After 5 h shaking at 37°C, OD was measured and pellets collected for plasmid prep.

### Sequencing Library Preparation

Library preparation used primers specific to flanking regions of the mutated region, with 5’ overhangs for Illumina indexing. First PCR (8 cycles) was followed by 1.0x SPRI bead clean-up, then indexed primer PCR (10 cycles) and 0.8x SPRI selection. Libraries were pooled equimolarly and sequenced on Illumina NovaSeq S2 (PE250) at the Wellcome Sanger DNA pipeline.

### Sequencing data processing

Raw read counts were processed with DiMSum v1.3.2^84^ using the following parameter settings: mutagenesisType = codon, fitnessMinInputCountAll = 10, vsearchMinQual = 20 and fitnessDropoutPseudocount = 0. For abundance assays, IC50 was estimated using the drc R package. However, IC50 fitting was limited by poor signal in dead mutants, reducing usable data. Instead, we selected three consecutive concentrations showing optimal separation between synonymous and nonsense variants, balancing resolution and noise. Scores were averaged across these biological replicates without weighting (as the shared inputs would penalise low-fitness variants due to Poisson error).

Abundance assay scores were normalised against the mode of synonymous and nonsense variants for each selected condition and then averaged. Binding assay scores were similarly normalised using the mode of synonymous and nonsense means, exploiting linearity across biological replicates.

### Structural analysis

All four TF-DBD in our assay bound to the corresponding motifs in the assay were modelled using AlphaFold3 ^65^, and compared to the closest solved structures in the PDB ^61,64,73,75,85^ database (Table 1). AlphaFold3 predictions were used as a reference for visualisation and analysis, including distance to DNA and rSASA calculation and pocket identification, to be consistent across all the TFs in our assay, since not all TFs have solved structures bound to DNA, nor all the mutated residues present in the solved structures. For instance, the structures of HNF1A bound to DNA are with different DNA-sequences and have some residues not solved or not included in the solved structures ^61,64^. There is no DNA-bound structure for FOXP1, and we used a highly conserved FOXP2-bound DNA (PDB:2a07) ^75^ as a proxy.

Core residues were defined as those with rSASA < 25%, DNA-contacting residues as those with ΔrSASA > 1% (bound vs unbound), and all others as surface residues. rSASA was computed using the built-in function in Pymol.

Residues defined as DNA-contacting by ΔrSASA included any putative DNA interface contacts to avoid bias in modelling the contribution of protein abundance to DNA binding.

Direct TF-DNA contacts were analysed and visualised using DNAproDB^60^. Euclidean distances to DNA were also calculated from the predicted DNA–protein complex using the ChimeraX “contacts” function with default parameters and a Van der Waals (VDW) overlap threshold of < 0.4 Å. All 3D Structure visualisation is all performed with ChimeraX-1.5.

Structural alignment of FOXG1 and FOXP1 DBD monomers was conducted using the ChimeraX “Matchmaker” function, employing the “Chain pairing” and “Best-aligning pair of chains” options. Sequence alignment was performed with the BLASTP “Blast 2 Sequences” tool ^86^.

Secondary structure annotation was performed using SSDraw^87^ on the AlphaFold3-predicted structure, with colouring based on the pLDDT confidence scores.

### Binding-residuals

To assess deviations from the expected relationship between abundance and binding, a LOESS regression model was fitted to predict binding scores as a function of abundance using mutations at positions that do not contact DNA. We used the loess () function using R^88^, set the Span parameter to 0.9, and used weights as the inverse variance of binding scores. Two focal points (0,0) and (−1, −1) were added with weights=1e+04, so that loess curves pass through the two focal points.

Residuals were calculated by subtracting the binding scores predicted by the LOESS model, based on the abundance scores, from the observed binding scores.

Per-position median absolute residuals were calculated by taking the median of all absolute residuals at each position, after excluding variants with very low abundance scores, which typically correspond to premature stop codons’ phenotypes (i.e., abundance < –0.8).

### Allosteric decay

We used the *nlsLM* function from the *minpack.lm* package in R to fit an exponential decay model to the absolute binding-residuals as a function of the minimum distance between each protein residue and DNA. The model followed the form: *y=a⋅exp(−b⋅x)+c*, where y represents the absolute residual and x the distance. Initial parameter values were set as follows: *a* was initialised to the maximum absolute residual, *c* to the minimum, and *b* to 0.5.

### Pocket prediction

Among the available tools, we selected PASSer (Protein Allosteric Sites Server) ^89,90^ due to its comprehensive integration of multiple algorithms, detailed reporting on the pockets, and capability for easy visualisation. Pocket identification and druggability prediction within PASSer is based on FPocket ^67^ druggability scores ranging from 0 to 1. 0 indicates unlikely to be bound by small molecules or peptides, and a score of 0.5 or above is generally considered a good, potentially druggable pocket..

We used the web server ensemble model, defining one Chain for the pocket identification by uploading the Alphafold3 predicted structure with all the mutated regions and the DNA included. Resulting pocket PDB files were visualised in ChimeraX.

### Clinical variant interpretation

Variant function classification was based on the following criteria: mutations were classified as neutral if the effect size was ≤ 0.1 and Benjamini-Hochberg FDR ≥ 0.05; as loss-of-function if the binding score was < −0.1 and FDR < 0.05; and as gain-of-function if the binding score was > 0.1 and FDR < 0.05. Variants with large effect sizes that were not statistically significant, or with small effect sizes that were statistically significant, were considered uncertain (Extended Data Fig. 6b).

Biophysical classification considered mutations as abundance-affecting if the residuals were < 0.1 (unless classified as neutral) and included “dead” variants with abundance scores below 80. Mutations were considered binding-affecting at the DNA interface if residuals were > 0.1, FDR < 0.05, and located at DNA-contacting positions (Extended Data Fig. 6c). Allosteric binding-affecting mutations were defined as those meeting the same criteria but located outside DNA-contacting positions (Extended Data Fig. 6c).

### Statistical tests

All statistical analyses were performed using R. To assess whether the functional scores (binding and abundance, respectively) of each variant significantly differ from zero across biological replicates, we performed one-sample t-tests. Resulting p-values were corrected for multiple testing using the Benjamini–Hochberg (BH) false discovery rate (FDR) method within each dataset.

To evaluate whether individual residue positions or specific groups of residues (e.g., core, surface, etc.) significantly differ from zero, we used one-sample Wilcoxon signed-rank tests. For datasets with fewer than 30 samples, Bonferroni correction was applied; for those with 30 or more samples, BH correction was used.

To compare variance between groups, we performed Kruskal–Wallis tests, followed by post hoc Dunn’s tests for pairwise comparisons.

## Data Availability

All DNA sequencing data have been deposited in the European Nucleotide Archive (ENA) at EMBL-EBI under accession number PRJEB97482

## Code Availability

Code is available via the public Github repository https://github.com/lehner-lab/TF-MAPS

## Acknowledgements

This study was funded by Wellcome (Grant reference: 220540/Z/20/A, ‘Wellcome Sanger Institute Quinquennial Review 2021-2026’). We thank Sanger DNA Pipelines team, especially Matthew Mayho for their assistance with Illumina Sequencing library preparation and sequencing services. We also thank Sanger HumGen/GenGen Informatics for their assistance with the computational infrastructure. We thank Jussi Taipale, and Lehner Team members for discussion and comments on the manuscript.

## Author Contributions

X.L. performed all experiments and analyses. X.L. and B.L. conceived the project, designed analyses, and wrote the manuscript.

---

**Extended Data Fig. 1:**
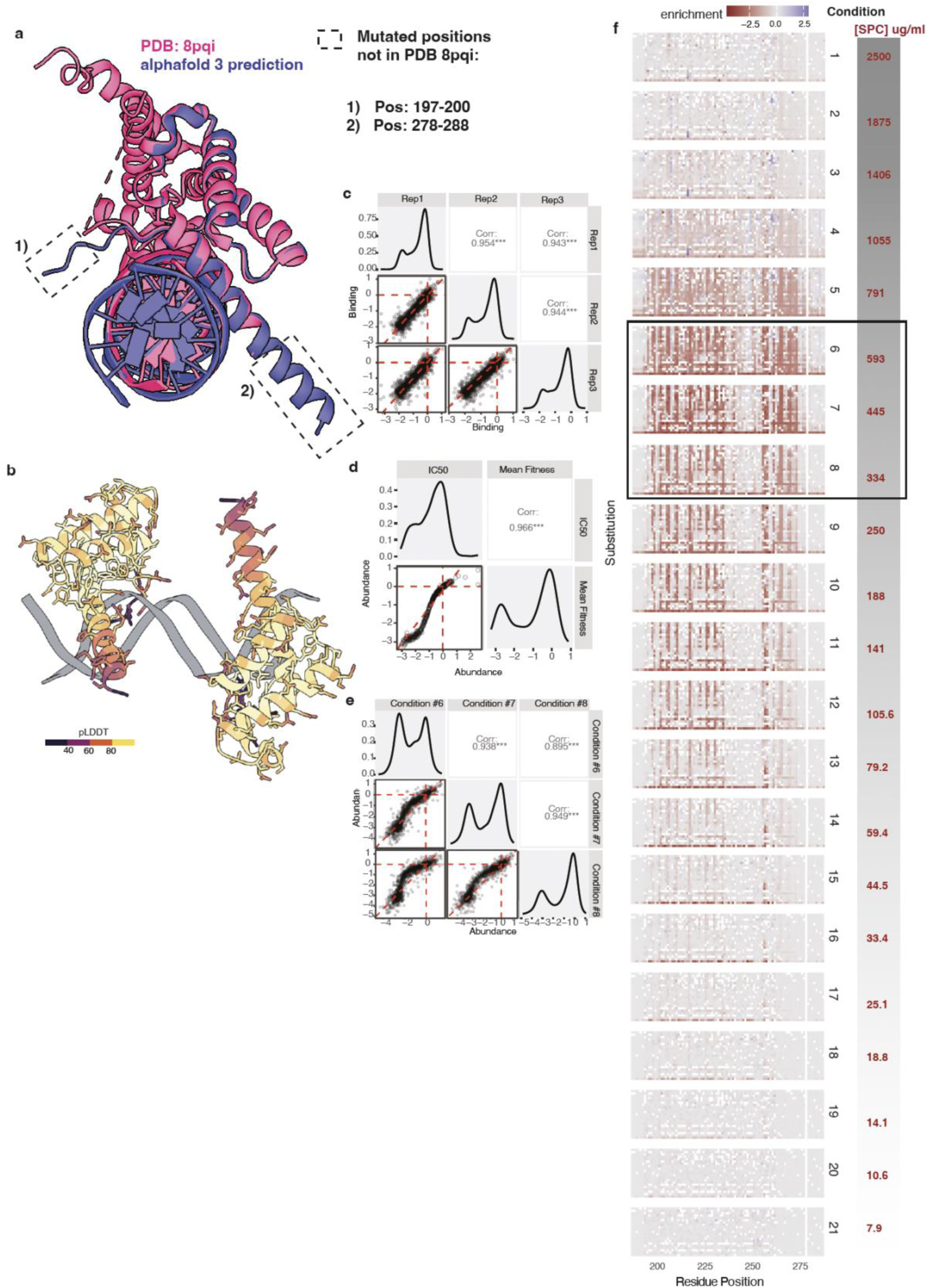
Reproducibility of the experiments with HNF1A POU_H_. (a) Overlay of the AlphaFold3-predicted DNA-bound HNF1A POUH region and the experimentally solved DNA-bound structure of the HNF1A DNA-binding domain. (b) AlphaFold3-predicted HNF1A POUH two monomers bound to the palindromic sequence of DHA. (c-e) Correlation matrices showing assay reproducibility for the binding assay (c) and abundance assay (d,e). Weighted means of abundance scores in (d) were taken from conditions #6, 7. and 8, which are shown individually in (e). (f) Heatmaps of the abundance assay across all 21 tested conditions with different concentrations of spectinomycin. Conditions #6,7,8 are highlighted.

**Extended Data Fig. 2:**
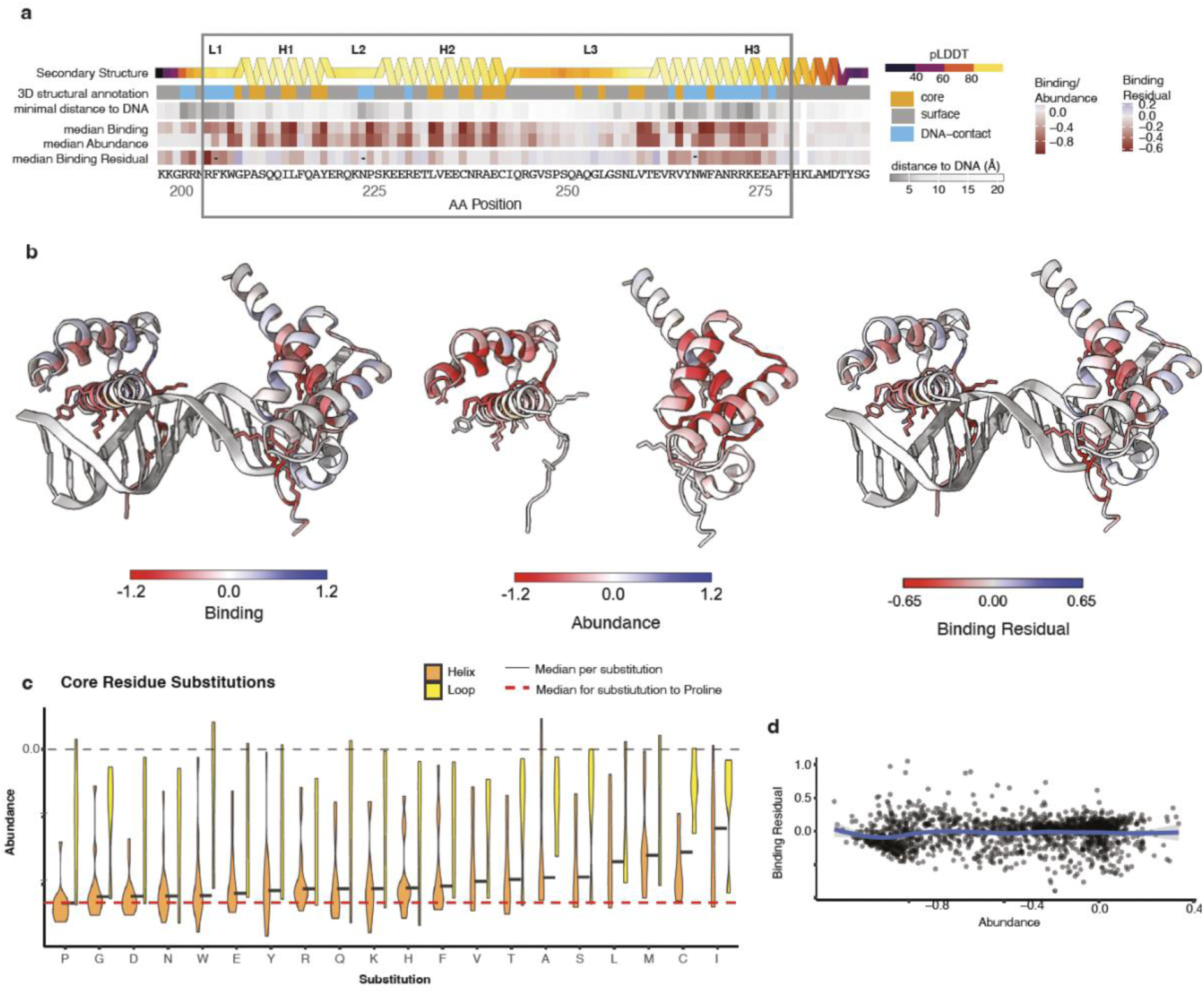
Distribution of HNF1A POU_H_ mutational effects on DNA binding and protein abundance. (a) Structural annotations and their positional median mutational effects, with the boxed region representing the conserved POUH annotation. (b) Structural view of the median mutational effects, with the side-chain atoms shown only for the DNA-contacting residues (b). (c) Comparison of abundance effects by substitutions to each amino acid. (d) Scatter plot comparing the relationship between binding-residuals and abundance for all missense mutations. Scatter plot showing the relationship between binding residuals and abundance for all missense mutations. The blue line represents the LOESS fit, and the grey shading indicates the corresponding 95% confidence interval.

**Extended Data Fig.3:**
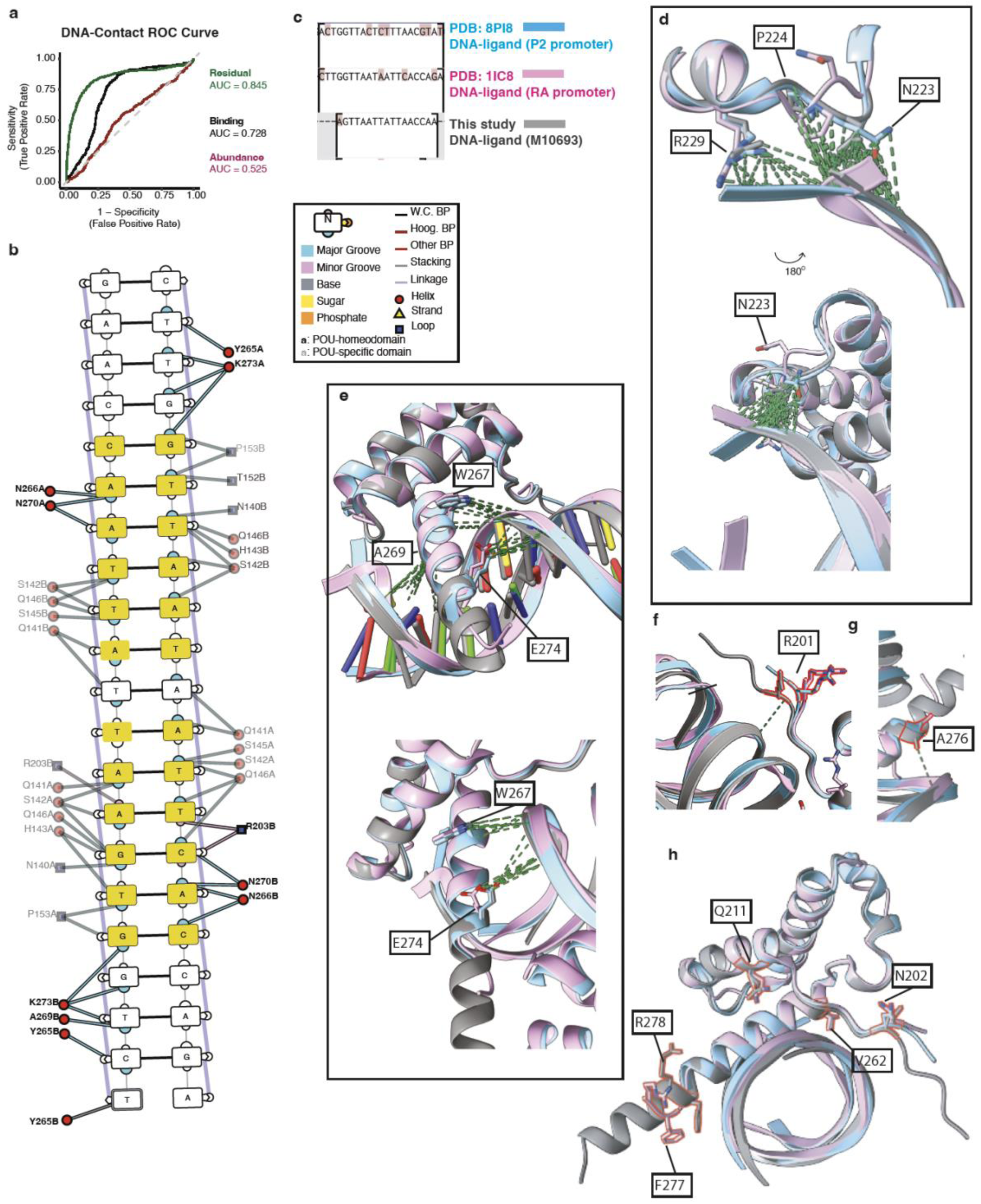
HNF1A POUH DNA-binding interface. (a) ROC curves for classifying DNA-contacting residues using positional median binding-residuals, as well as binding and abundance scores. (b) Illustration of DNA-contacting residues based on PDB 1ICS. (c) Sequence alignment of DNA used in this study and in the two published structures. Non-identical bases are highlighted. (e-h) Three aligned structures of HNF1A POUH bound to DNA, viewed from different angles highlighting different positions.

**Extended Data Fig. 4:**
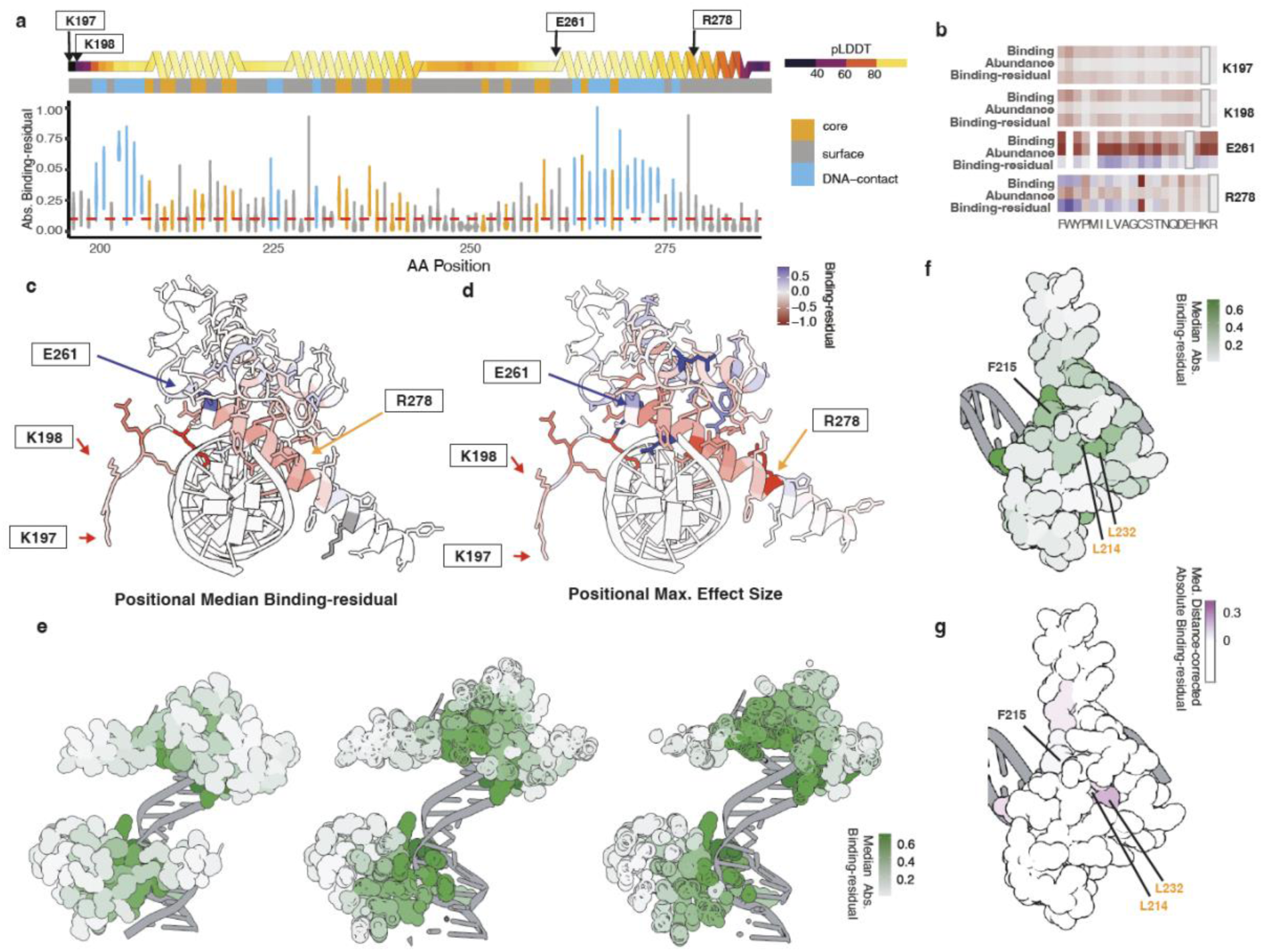
DNA-binding-residuals for HNF1A POU_H_. (a) Absolute binding-residuals plotted along the amino acid sequence of HNF1A POUH. (b-d) Examples of surface-exposed positions showing large absolute residuals for all mutations (b), highlighted on positional medians (c) and maximum effect sizes (d), illustrated on 30 structural views. (e) Structural view of two HNF1A POUH monomers bound to DNA, with absolute residuals colour-coded by magnitude. (f, g) Structural view of the HNF1A POUH monomer bound to DNA , shown from the side opposite the DNA. The structures are coloured by the positional median absolute binding-residuals(f), and residuals after accounting for the distance-dependent decay (g). Positions with higher-than­ expected distance-corrected binding residuals are highlighted.

**Extended Data Fig. 5:**
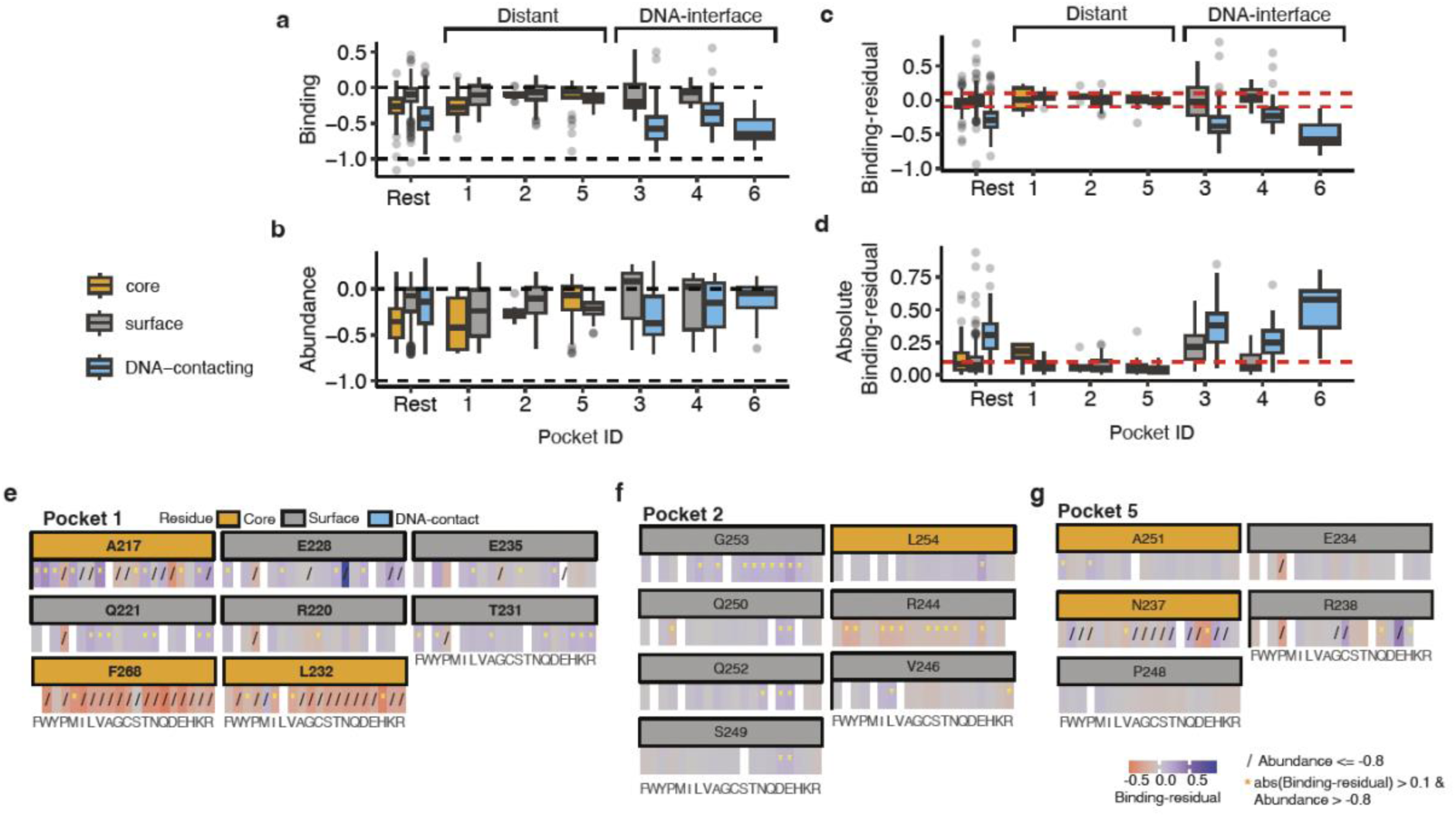
HNF1A POU_H_ pockets. (a-d) Comparison of mutational effects for residues within each pocket versus those outside the pockets in HNF1A POU_H_. Mutations with abundance scores below -0.8 are excluded. Red dotted lines denote an effect size threshold of 0.1. (e-g) Binding-residuals of all mutations located at distal pockets: in Pocket 1 (e), Pocket 2 (f), and Pocket 5 (g).

**Extended Data Fig. 6:**
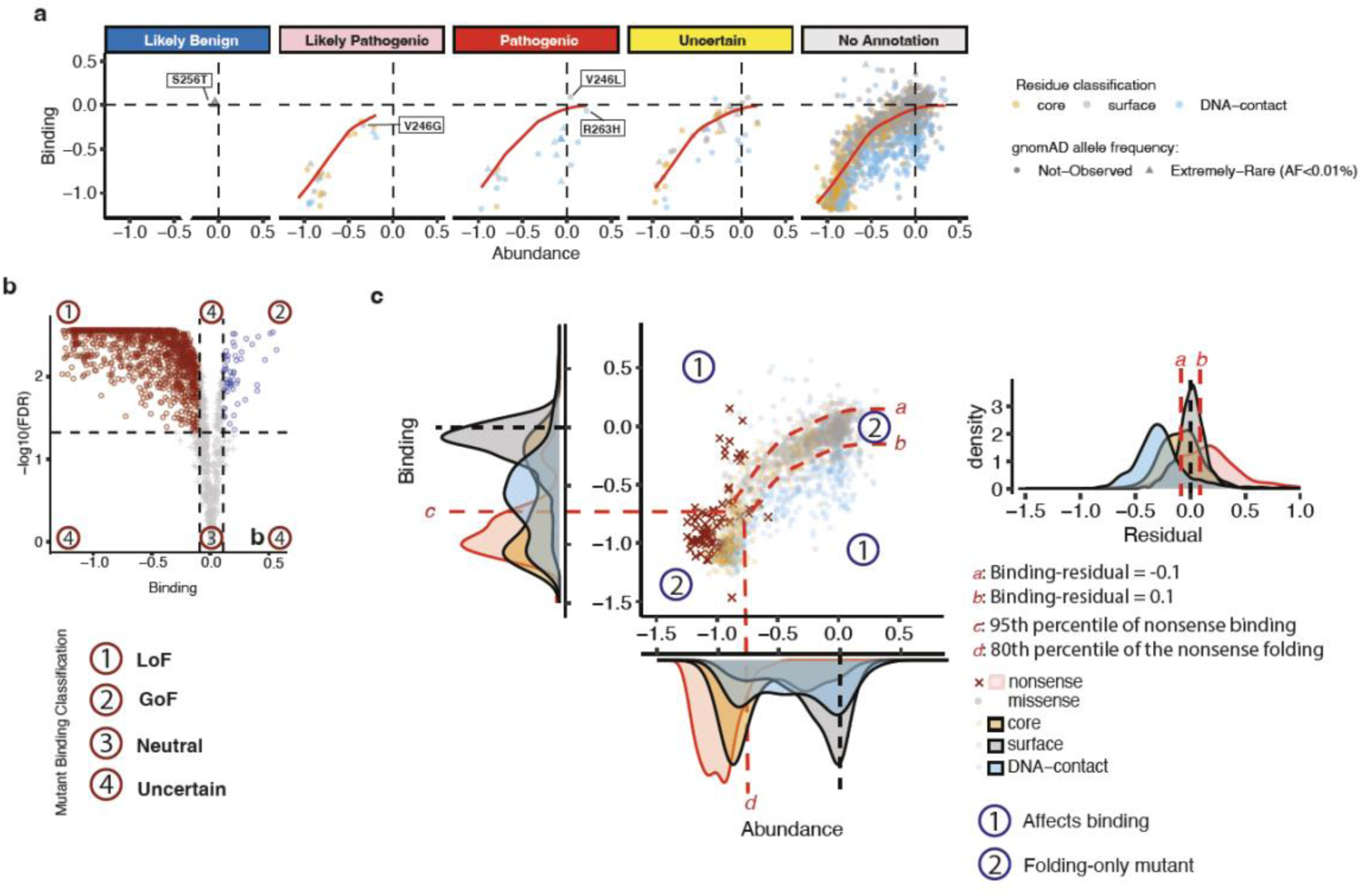
Classification of the functional effects of HNF1A POU_H_ variants. (a) Clinical and non-annotated variants mapped in the Abundance-Binding phenotype space. (b, c) Strategy for classifying variants by functional impacts, based on the volcano plot of binding (b), and by biophysical mechanism based on the Abundance-Binding scatter plot (c).

**Extended Data Fig. 7:**
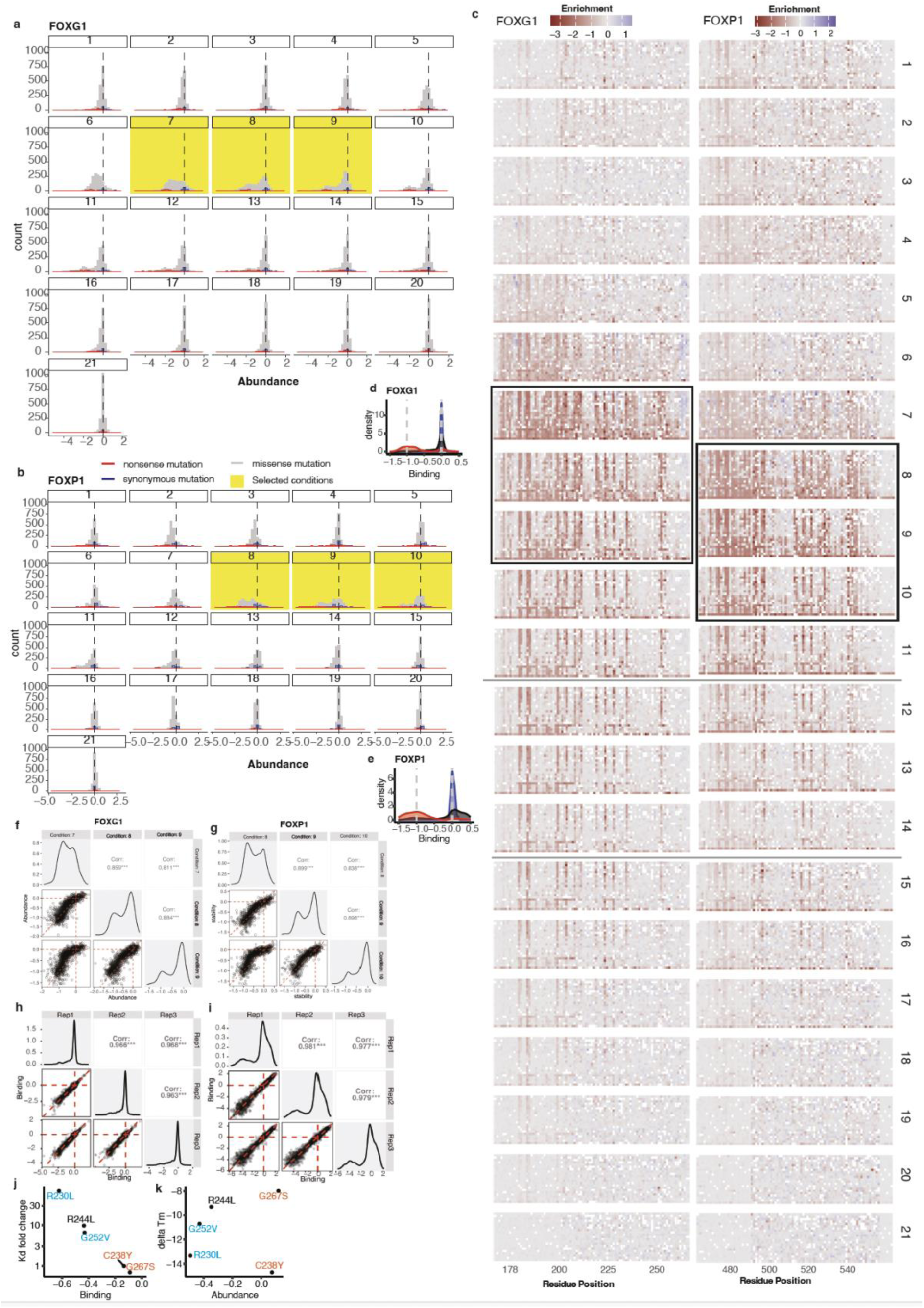
Reproducibility of the experiments with FOXP1 and FOXG1 DBDs. (a-c) Density plots (a, b) and heatmaps (c) of abundance assays across all 21 tested conditions with varying concentrations of spectinomycin, where condition “1” is the highest concentration and “21” the lowest for FOXG1 (a) and FOXP1 (b). Three highlighted conditions in each panel were selected for calculating abundance scores. (d, e) Density plots of binding scores after normalisation across three biological replicates for FOXG1 (d) and FOXP1(e). (f-i) Correlation matrices showing assay reproducibility for the abundance assays (f, g) and binding assays (h, i) for FOXG1 (f,h) and FOXP1 (g,i). (j, k) Correlation between published Kd values and FOXG1 binding scores (j), Tm changes and abundance scores (k). Each mutation is coloured according to the structural annotation of its residue: cyan for DNA-binding interface residues, pink for protein core residues, and black for surface residues.

**Extended Data Flg. 8:**
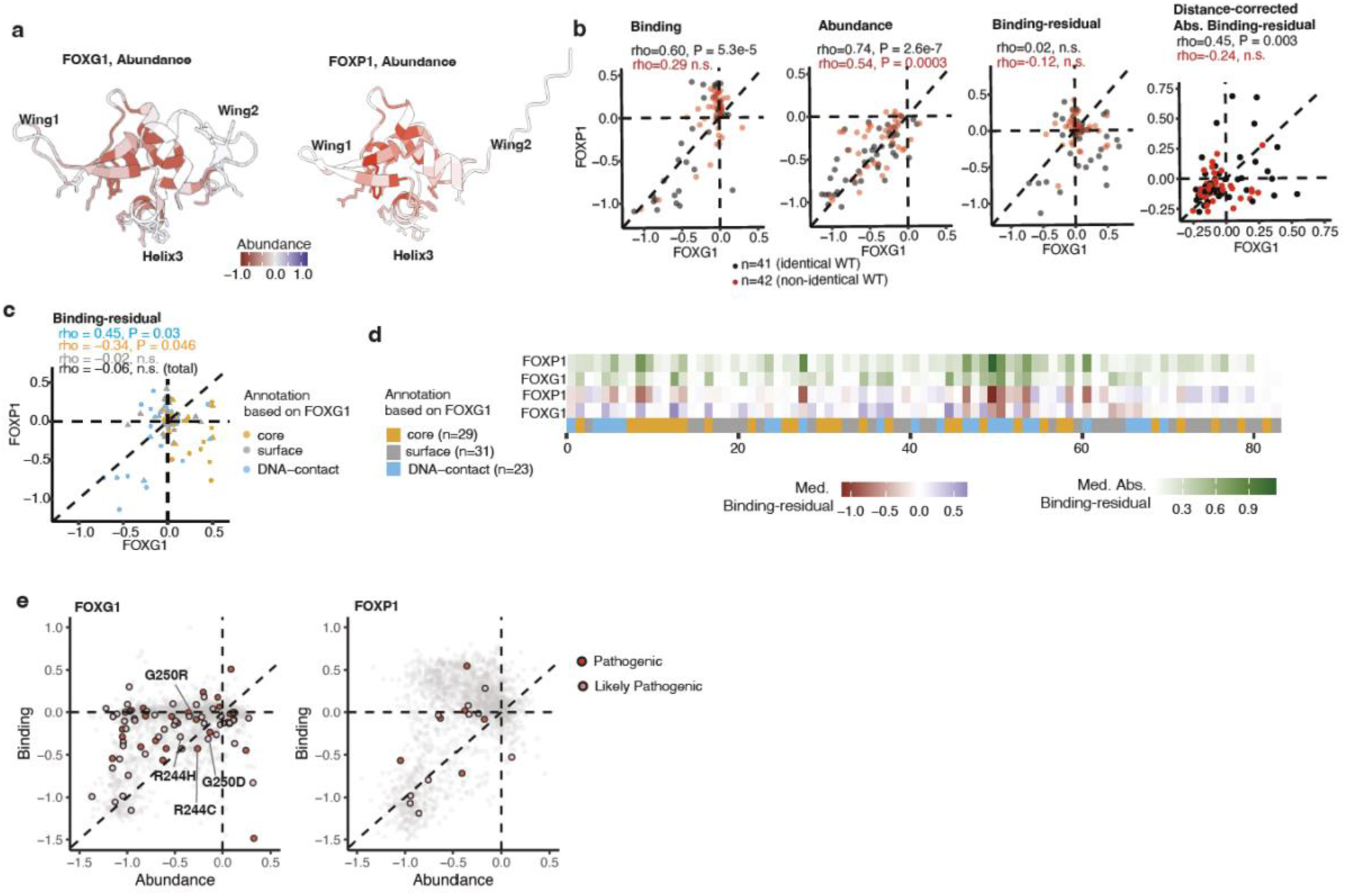
TF-MAPS positional median comparisons for FOXG1 and FOXP1 forkhead family DBDs. (a) Median mutational effects per amino acid residue for FOXG1 abundance. (b) Correlations between matching positions in FOXG1 and FOXP1 for median binding, abundance, binding-residuals, and distance-corrected absolute binding-residual, colour­ coded by identical and non-identical matched positions. (c) Correlations between median binding residuals of matched positions in FOXG1 and FOXP1 coloured by structural annotation. (d) Heatmaps of aligned positional median binding­ residuals and absolute binding-residuals for FOXG1 and FOXP1. Positions are re-indexed based on sequence alignment. (e) Two-dimensional binding-abundance landscape highlighting clinically annotated pathogenic and likely pathogenic variants.

**Extended Data Flg. 9:**
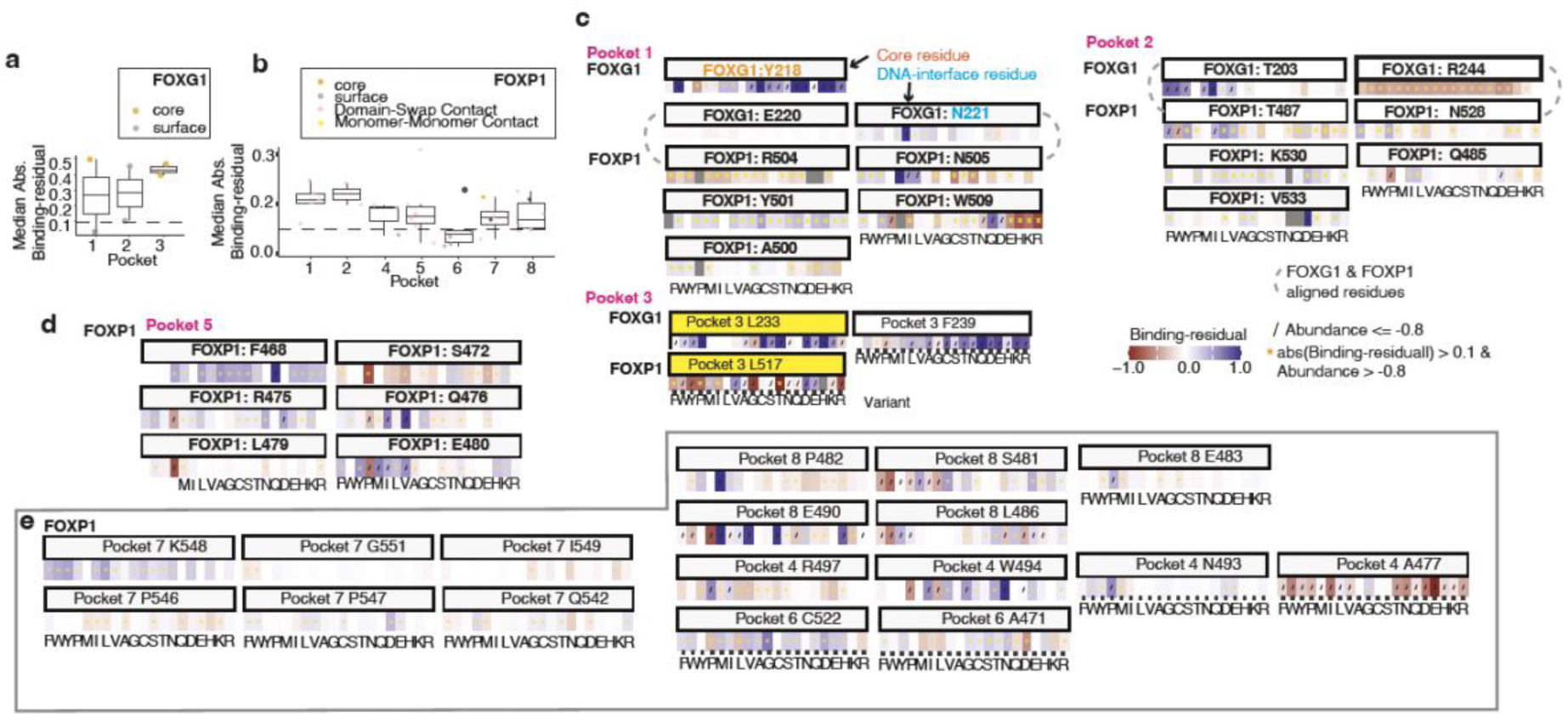
TF-MAPS positional median comparisons for FOXG1 and FOXP1 for1<head family DBDs. (a, b) Median absolute binding-residuals of each residue within each pocket of FOXG1 (a) and FOXP1 (b). Only residue outside the DNA interface, and with abundance score> -0.8 are shown. Pocket 3 of FOXP1 is not shown because the only non-DNA contacting residue L517 is highly destabilising with the median abundance score <-0.8. (c) Binding-residuals of aligned pockets of FOXG1 and FOXP1. DNA-contacting positions are excluded except for a DNA-contacting FOXG1 residue N221 of Pocket 1, to compare with its aligned non-DNA-contacting FOXP1 residue N505. (d, e) Binding residuals of Pocket 5 of FOXP1 (d) and other FOXP1 pockets (e).

**Supplementary Table 1.**
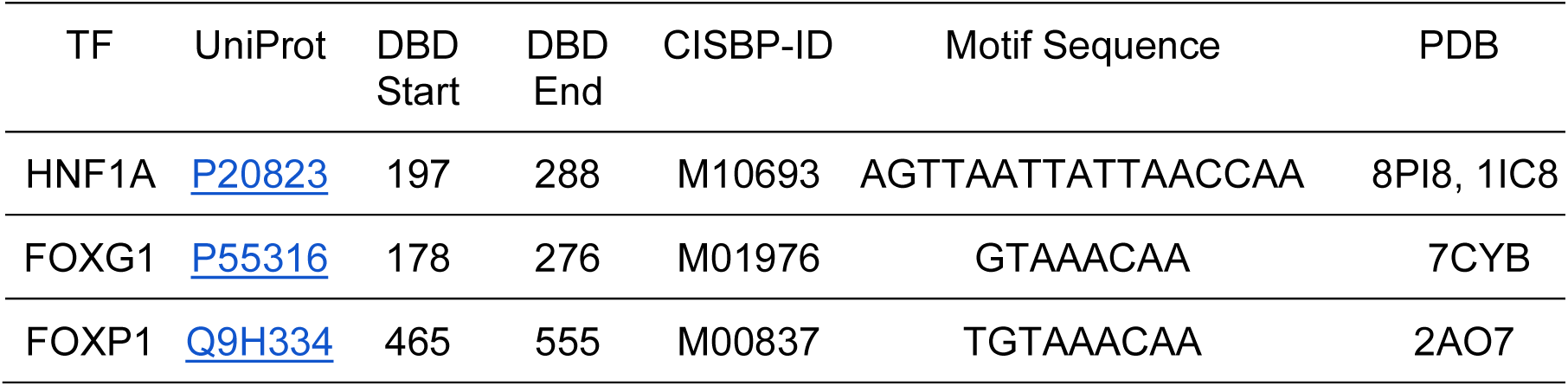
DBDs and Motifs.

**Supplementary Table 2.** HNF1A POU_H_ Pockets.

| Pocket | Type | Sum of Absolute Binding-residuals | Druggability |
| --- | --- | --- | --- |
| 1 | **Distant** | 1.79 | 0.247 |
| 2 | **Distant** | 0.62 | 0.108 |
| 3 | Interface | 0.43 | 0.003 |
| 4 | Interface | 0.56 | 0.001 |
| 5 | **Distant** | 0.19 | 0.001 |
| 6 | Interface | 1.01 | 0.000 |

**Supplementary Table 3.** FOXG1 FKH Pockets.

| Pocket | Type | Sum of Absolute Binding-residuals | Druggability |
| --- | --- | --- | --- |
| 1 | Interface | 0.44 | 0.420 |
| 2 | Interface | 1.79 | 0.070 |
| 3 | Interface | 0.83 | 0.000 |

**Supplementary Table 4.** FOXP1 FKH Pockets.

| Pocket | Type | Sum of Absolute Binding-residuals | Druggability |
| --- | --- | --- | --- |
| 1 | Interface | 1.83 | 0.494 |
| 2 | Interface | 0.57 | 0.124 |
| 3 | Interface | 1.81 | 0.048 |
| 4 | Interface | 0.45 | 0.041 |
| 5 | **Distant** | 1.39 | 0.013 |
| 6 | Interface | 3.04 | 0.007 |
| 7 | **Distant** | 1.14 | 0.002 |
| 8 | **Distant** | 0.57 | 0.001 |

